# Near-critical tuning of cooperativity revealed by spontaneous switching in a protein signalling array

**DOI:** 10.1101/2022.12.04.518992

**Authors:** Johannes M Keegstra, Fotios Avgidis, Yuval Mulla, John S Parkinson, Thomas S Shimizu

## Abstract

Dynamic properties of allosteric complexes are crucial for cellular information processing. However, direct observations of allosteric switches have been limited to compact molecular assemblies. Here, we report *in vivo* FRET measurements of spontaneous discrete-level fluctuations in the activity of the *Escherichia coli* chemosensory array — an extensive membrane-associated assembly comprising thousands of molecules. Finite-size scaling analysis of the temporal statistics by a two-dimensional conformational spread model revealed nearest-neighbor coupling strengths within 3% of the Ising second-order phase transition, indicating that chemosensory arrays are poised at criticality. Our analysis yields estimates for the intrinsic timescale of conformational changes (~ 10 ms) of allosteric units, and identifies near-critical tuning as a design principle for balancing the inherent tradeoff between response amplitude and response speed in higher-order signaling assemblies.

**One-setence summary:** *In vivo* measurements of protein signaling array fluctuations reveal an allosteric system poised at criticality.

## Main Text

Large protein assemblies at the heart of many cell signaling processes exhibit varying degrees of structural and dynamical order, from liquid-like granules (*1, 2*) to solid-like arrays (*3–5*). Elucidating the mechanisms and design principles of signaling assemblies would not only advance our understanding of cell signaling in nature, but also boost current efforts towards rational design of synthetic protein circuits (*6, 7*). Recent experiments are revealing how phase transitions leading to the formation of liquid-like assemblies — first-order transitions analogous to the condensation of gas into liquid — are used by cells to implement various information-processing tasks, such as signal initiation and confinement (*8*), kinetic proofreading (*9*), and noise control (*10*). Theory also predicts that a different phase transition can occur within solid-like assemblies — a second-order transition arising from conformational interactions between protein subunits (*11*). However, requirements for second-order phase transitions are more stringent than those for first-order transitions — they occur only at a special point in phase space, a critical point. It remains unclear if such second-order transitions occur in signalling assemblies and if yes, how they constrain the functional design of protein assemblies.

Recent structural studies are revealing an increasing number of solid-like signaling assemblies with a high degree of spatial order, where subunit proteins are arrayed in a regular pattern. The repertoire of such crystal-like assemblies reported to date is diverse in both form and function, and includes one-dimensional filament-like assemblies found in cellular homeostasis and inflammation signaling (*12, 13*), protein rings that mediate control of cell motility and apoptosis (*14, 15*), as well as two-dimensional arrays involved in chemosensing, neuromuscular control, and innate immune responses (*16–19*). Yet, despite exciting advances in resolving the ultrastructure of these large assemblies, mechanistic design principles of their signaling function remain challenging to address experimentally. By contrast to compact oligomeric signaling complexes of fixed size, these extensive structures tend to assemble through open-ended polymerization and/or multivalent interactions, making them inherently variable in both size and composition (*4*). The resulting polydispersity and stoichiometric diversity tend to mask their true signaling dynamics, as well as their size- and composition-dependence, both *in vivo* and *in vitro.* An ideal functional assay would therefore interrogate assembly-level dynamics *in singulo* — at the level of an individual assembly — but experimental realization has remained elusive.

Here we report *in vivo* FRET experiments that resolve this challenge for the chemosensory array of *Escherichia coli,* a canonical two-dimensional signaling assembly lining the cytoplasmic membrane, which allows motile bacteria to navigate chemical environments with exquisite sensitivity (*20–22*). This higher-order assembly comprises thousands of receptor, kinase, and scaffolding molecules arranged in a well-defined lattice structure (*23–25*) and thus serves as a paradigm for signaling in two-dimensional protein assemblies. The *in vivo* signaling activity of chemosensory arrays can be read out by an intermolecular FRET system (*20*) that reports on CheA kinase activity, the signaling output of arrays, by labeling two downstream proteins: the response regulator CheY, which receives phosphoryl groups from CheA, and its phosphatase CheZ (Fig. 1a). Our strategy leverages recent advances in extending this *in vivo* FRET technique to the single-cell level (*26, 27*), to achieve functional measurements of individual arrays within live cells. Imaging studies have revealed that the partitioning of chemoreceptors and associated proteins into arrays is highly variable across cells, even within isogenic populations, with some cells possessing many small arrays while others have only one or a few large arrays (*28, 29*). Thus, by searching for cells in which signaling is dominated by a single large array, we aimed to achieve *in singulo* measurements of functional array output.

**Figure 1:**
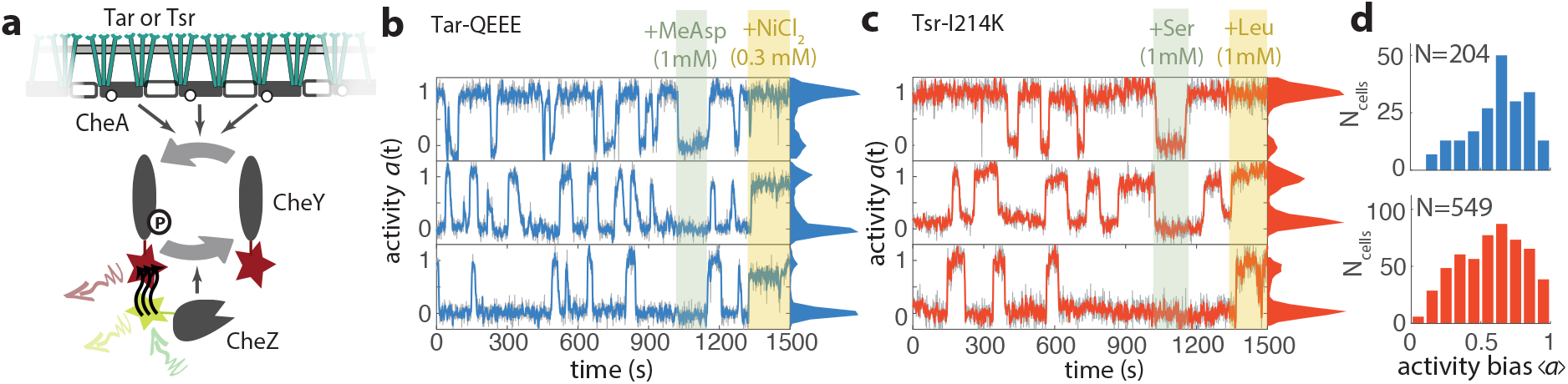
Chemosensory arrays of both major receptor types Tar and Tsr demonstrate switching fluctuations in the absence of sensory stimulation. **(a)**. Illustration of single-cell FRET assay to measure the activity of the CheA kinase embedded in an array of chemoreceptors (Tar or Tsr) (*26*). Measured FRET signal between fluorescently labeled CheZ and CheY is proportional to the activity of the chemosensory array. **(b)** Time series (raw in grey, 3s-averaged in blue) and histograms (on right margin) of activity determined by FRET, in three representative cells expressing only the chemoreceptor Tar [QEEE] without adaptation enzymes. Cells are exposed to measurement buffer (MotM) for most of the experiment and when indicated to an attractant stimulus of 1 mM *α*–methylaspartate (MeAsp) and a repellent stimulus of 0.3 mM NiCl_2_ to obtain the minimum and maximum FRET levels, respectively, that were then used to normalize the activity time series of each cell between zero and unity. **(c)** As in (b) but with cells expressing Tsr-I214K and 1 mM Serine (Ser) and 1 mM Leucine (Leu) used as attractant and repellent stimuli, respectively. **(d)** Histogram of activity bias in the absence of chemoeffectors 〈*a*〉, for Tar [QEEE] (top) and Tsr-I214K (bottom).

To identify cells whose chemoreceptor population is concentrated into a single large array, we focused on spontaneous fluctuations in kinase output that are observable by FRET at the single-cell level. Previous single-cell FRET measurements revealed substantial spontaneous fluctuations of chemoreceptor array activity in the absence of changes in external ligand stimuli or covalent modification feedback from downstream adaptation enzymes (*26, 27*). The amplitude of these residual fluctuations was substantially augmented in cells engineered to express only a single chemoreceptor species (out of five in wildtype *E. coli*), and, strikingly, a subset of these cells demonstrated spontaneous two-level switching between fully active and inactive states while the input ligand stimulus was held constant (*26*). Spontaneous two-state switching is a hallmark of allosteric signaling complexes such as ion channels (*30*), but their observation requires measurements at the level of individual complexes; ensemble-averaged experiments can not resolve the switching dynamics because the timing of switching events are uncorrelated and thus the switching dynamics are averaged out. Because in our FRET experiment the readout integrates the kinase output of all arrays within a cell, a plausible explanation for the two-level switching fluctuations observed in FRET is that they reflect stochastic switching in the signaling state of one large dominant array. However, it is also possible that other factors, such as spurious fluctuations in ligand signals, are driving synchronous switching of a larger number of otherwise independent signaling arrays.

To gain better insight into the mechanism that generates two-level switching, we measured temporal fluctuations by single-cell FRET recordings in thousands of individual cells. We first confirmed robust two-level switching in the FRET signal over time within cells expressing the aspartate receptor Tar as the sole chemoreceptor in a genetic background devoid of the adaptation enzymes CheR and CheB (Fig. 1b). Wild-type Tar in the absence of adaptation enzymes has a steady-state activity 〈*a*〉 (defined as the average activity bias in the absence of chemoeffectors) close to unity, and does not switch unless this bias is shifted down by chemoeffector ligands (not shown). To rule out exogenous ligand fluctuations as the driver of chemoreceptor array switching, in these experiments we instead tuned down the activity bias by expressing Tar in the QEEE covalent modification state by a single-residue replacement at one of four adaptation-modification residues (from QEQE in wildtype Tar), which yields an intermediate activity bias without the addition of chemoeffectors (see Supplementary text, Fig. 1b,d) (*31*). In 204 of 1414 cells obtained from 19 FRET experiments, we observed switching between a high and low activity state (> 65% of transitions showing activity level changes of > 0.7). Spontaneous switching in the absence of exogenous ligands was not specific to Tar receptors, as we observed similar switching behavior (two-state switching in 548/4446 cells, across 44 experiments) in analogous experiments with Tsr-I214K, a single-residue replacement mutant of the serine receptor Tsr within its ‘control cable’ region (*32*) with a down-shifted activity bias similar to that of Tar [QEEE] (Fig 1c). To rule out that switching fluctuations were driven by ligands secreted by the cells themselves, we also tested single-residue replacement mutants of the ligand binding pocket (residue 69) of both Tar [QEEE] and Tsr-I214K known to abolish ligand binding in both receptor types (*33, 34*). Cells expressing these mutated receptors showed no response to their cognate ligands, as expected, but still exhibited switching behavior (Fig. S2). Taken together, these observations suggest that two-state switching reflects a generic property of chemosensory arrays that is not specific to a single chemoreceptor species, and that it is driven by intrinsic fluctuations within the array, rather than extrinsic fluctuations of ligand inputs.

Two-level switching in whole-cell kinase activity is surprising because it implies synchronous switching of very large molecular populations — *E. coli* chemoreceptors and their associated kinases are expressed at a level of ~ 10^3^ – 10^4^ copies per cell (*35*). We ruled out that switching cells have an anomalously low expression of chemoreceptor array components by confirming that the maximum amplitude of the CheY-CheZ FRET signal, which is known to be proportional to the amount of receptor-kinase complex (*20, 26*), was comparable between switching and non-switching cells (Fig. S1e). Furthermore, the fraction of cells in which only a single array was visible by fluorescence microscopy (13%, Fig. S1e) was very similar to the fraction of cells demonstrating two-state switching (14% for Tar and 12% for Tsr, Fig. S1c). Finally, cells expressing Tsr-F396Y, a single-residue replacement mutant defective in response cooperativity, demonstrated no switching behavior (Fig. S3), indicating that the switching phenotype is linked to cooperativity within the array. These observations add further support to the idea that the two-level fluctuations observed in the FRET signal result from cooperative switching of thousands of molecules within a single large chemosensory array.

To consider the physical mechanism underlying this long-range cooperativity, we further dissected the temporal statistics of switching fluctuations (Fig 2a) by extracting the time interval between switching events, hereafter called the residence time Δ*t*_up_ and Δ*t*_down_ for time spent in the up (*a* ≈ 1) and down (*a* ≈ 0) state, respectively, as well as the duration of the activity transient upon switching, hereafter called the transition time *τ*_+_ and *τ*_-_ for upward and downward switches, respectively. We first interpreted these data as a barrier-crossing stochastic process in which the up and down states correspond to wells within an energy landscape (Fig 2b), the shape of which could be approximated from the observed activity time series histograms with the free energy difference *ΔG* (in units of the thermal energy *k_B_T*) between the up and down states determined by the activity bias 〈*a*〉 as Δ*G* = ln[(1 – 〈*a*〉)/ 〈*a*〉]. Consistent with this picture, we found that for both Tar [QEEE] and Tsr-I214K arrays, the residence times were exponentially distributed across the full range of 〈*a*〉 (Fig. 2c), and the average residence time of each cell 〈Δ*t*_up/down_〈 as a function of Δ*G* was found to obey an Arrhenius-type exponential scaling 〈Δ*t*_up/down_〉 (Δ*G*) = *τ_r_*e^−*γ*^_up/down_^Δ*G*^ (Fig. 2d), where *τ_r_* is a characteristic residence timescale and *γ*_up/down_ are fitting constants. Thus, at the level of residence-time statistics, both Tar and Tsr arrays behave like a Brownian particle diffusing in a double-well potential along a “reaction coordinate” corresponding to the array’s activity *a* (*36*). From the crossings of the Arrhenius fit lines in Fig. 2d, we determined *τ_r_* for both receptor species: 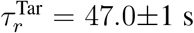 and 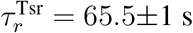.

**Figure 2:**
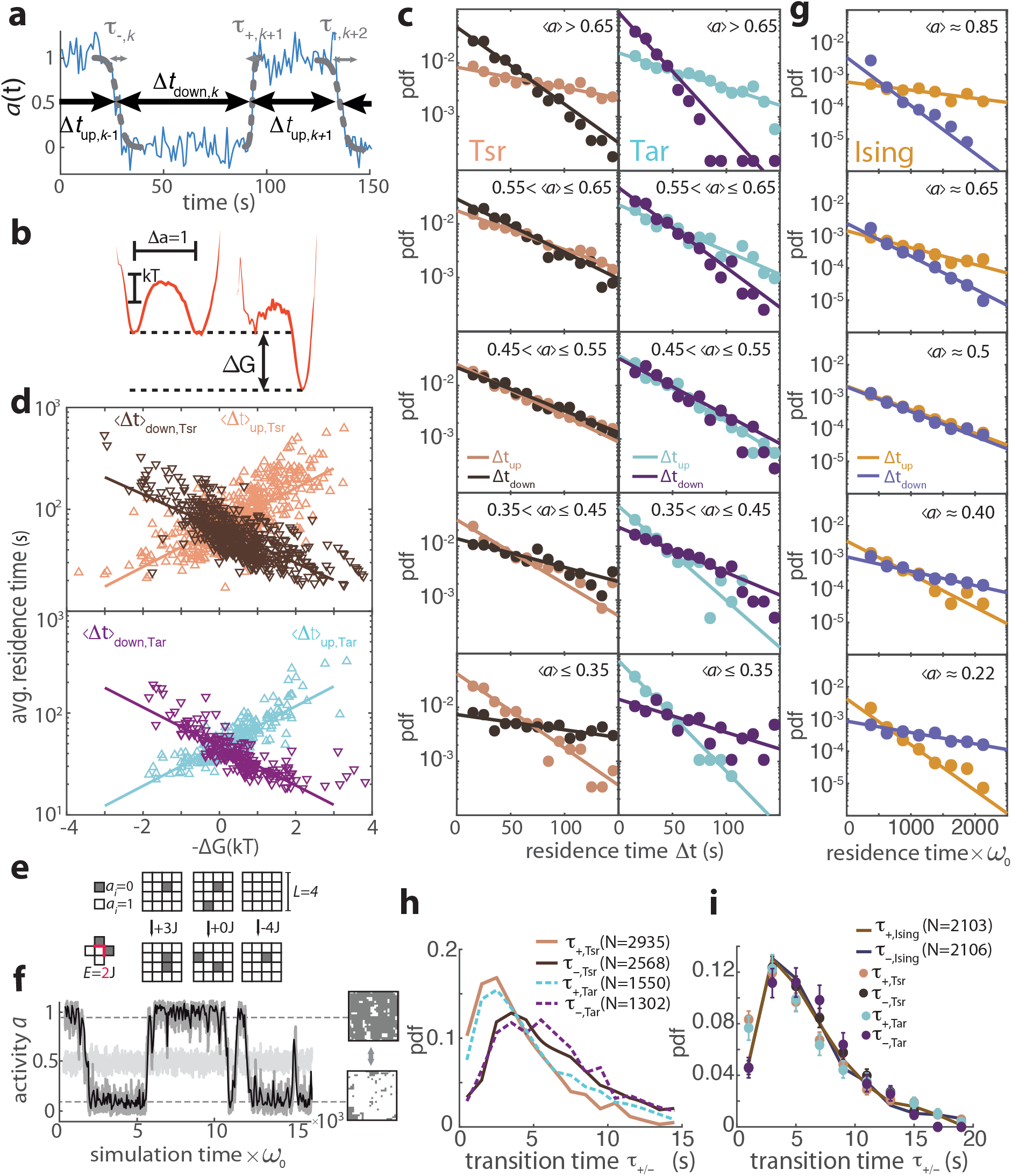
Temporal statistics of switching events are well described by an Ising-type conformational spread model. **(a)** Definitions of residence times Δ*t*_up/down_ and transition times *τ*_+/-_, determined routinely for each switching event (see Materials and Methods). **(b)** Coarse-grained energy landscape along the array-activity coordiate a (estimated as the negative logarithm of the activity histogram) based on selected time series with 〈*a*〉 ≈ 0.5 (15 cells, left) and 〈*a*〉 ≈ 0.9 (12 cells, right) from cells expressing Tsr-I214K. **(c)** Histogram of residence times from experiments. Residence times for Tsr-I214K (left) and Tar [QEEE] (right), with each event sorted by activity bias of the corresponding cell (colors as in panel d). In each panel, data (points) are shown together with fits to single exponential functions (solid lines). Fit parameters and number of data points are shown in Tables S2 and S4. **(d)** Mean residence times per cell as a function of the energy bias ΔG between the high and low activity state for all cells expressing Tsr-I214K (top) and Tar [QEEE] (bottom). Fit parameters and number of data points are shown in Table S6. **(e)** Conformational spread model of chemosensory array activity with size *L* × *L*, in which each individual unit can switch between activity states of 1 (white) or 0 (dark). A difference in neighboring spins is associated with an energy cost of *J*, shown for three different transitions on a lattice with size *L* = 4 in the absence of external field (*H* = 0). **(f)** Example activity time series obtained by simulating dynamics on a strongly coupled (*L* = 26, *J* = 0.4625kT, dark grey) and weakly coupled lattice (*L* = 20, *J* = 0.2375kT, light gray). The strongly coupled lattice exhibits stochastic switches between two activity levels (dashed lines), with representative simulated array states for each activity level (right). The simulated time series was downscaled to approximate the experimental acquisition frequency (solid black line). **(g)** As in panel (d), but histogram of residence times from simulated time series with *L* = 12 and *J* = 0.5kT, sorted by the activity bias generated by an applied indicated external field *H*. Fit parameters and number of data points are shown in Table S9. **(h)** Histogram of switching events for Tsr-I214K (solid lines) and Tar [QEEE] (dashed lines). **(i)** Transition times from simulated two-state time series (*N* = 12, *J* = 0.5kT, varying *H*). To approximate the experimental signal-to-noise ratio, gaussian white noise was added to the simulated time series (see Fig. S8). Experimental (points) and simulated (solid lines) histograms are scaled horizontally to have mean transition time of cells expressing Tsr for comparison.

To investigate whether and how the observed temporal statistics could be explained by allosteric subunit interactions, we turned to theory. We used an Ising-type conformational spread model of allostery (*37–39*), which assumes that subunit conformations are coupled through nearest-neighbor interactions. The strength of these interactions is parameterized by a coupling energy *J* (promoting order) whose magnitude relative to *k_B_T* (promoting disorder) determines a finite spatial range (*i.e.* a correlation length) over which action at one site can affect distant sites. By varying *J*, therefore, the model can represent allosteric systems along a continuous scale of conformational disorder, including that of the more widely used Monod-Wyman-Changeux (MWC) model (which is recovered upon taking the fully ordered limit *J* → ∞). Importantly for signaling function, two-dimensional Ising models are known to exhibit a second-order phase transition as a function of *J*, with a spontaneous ordering of subunit conformations above a critical coupling energy *J*\*. Although, to date, strong experimental support for conformational spread models has been obtained in one-dimensional protein rings (*40, 41*), Ising models exhibit a critical point only in two or higher dimensions, and the implications of the Ising second-order transition in two-dimensional protein assemblies remain untested experimentally.

We performed kinetic Monte Carlo simulations on an *L* × *L* lattice of allosteric units, each of which can flip between two conformational states, active (*a* = 1) or inactive (*a* = 0) (Fig. 2e and Materials and Methods). The flipping rate of the unit at site *i* was modified from a fundamental flipping frequency *ω*_0_ by the influence of its nearest neighbors (*j* ∈ 1,…, *N_j_*, where *N_j_* is the number of nearest neighbors) through the coupling energy *J* (in units of *k_B_T*) as *ω* = *ω*_0_*e*^-*J*(1–2a_*i*_) ∑_*j*_(1–2*a_j_*)^, corresponding to an Ising model in which each active-inactive bond on the lattice contributes an energy penalty of *J* (see Materials and Methods). At low coupling strength (*J* ≪ *J*\*), each unit switches independently and the total activity of the array demonstrates only small fluctuations about its mean value at 〈*a*〉 = 1/2 (Fig. 2e), but as the coupling energy is increased toward its critical value *J*\*, the correlation length exceeds the finite size of the array, which then begins to switch between fully active and fully inactive states (Fig. 2f, Fig. S7). We analyzed the temporal activity statistics of a simulated array with parameters within this two-state switching regime (*J* = 0.5 *k_B_T*, *L* = 12), with various values of a weak biasing field *H_b_* that modifies the flipping rate by a factor *e*^*H_b_*(1–2*a*)^ (see Materials and Methods), to approximate the diverse FRET activity biases observed across individual cells in the population (Fig.1c). Simulated residence time distributions (Fig. 2g) were in excellent agreement with their experimental counterparts (Fig. 2c), recapitulating at each activity bias their characteristic exponential shape.

By contrast, the measured transition time distributions had a peaked profile for both Tar and Tsr arrays (Fig. 2h). The average downward transition time 〈*τ*_-_〉 and the average upward transition time 〈*τ*_+_〉 were similar between Tar and Tsr arrays, with 〈*τ*_-_〉 slightly greather than 〈*τ*_+_〉 in both cases (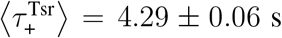, 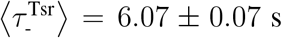, 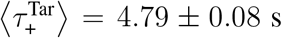 and 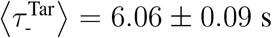; mean ± s.e.m.). Remarkably, however, when normalized by these mean values, all measured time distributions collapsed onto a common profile that, in turn, was in excellent agreement with its simulated counterparts (Fig. 2i). Furthermore, both measured and simulated transition-time statistics demonstrated no dependency on the activity bias (Fig. S5 and S9). Collectively, the high degree of quantitative agreement between these measured and simulated temporal statistics suggest that an Ising-type conformational spread model with a near-critical coupling strength (*J* ≈ *J*\*) provides an excellent approximation to chemoreceptor array dynamics.

We sought to quantify how close to criticality both Tar and Tsr arrays are tuned. The behavior of Ising-type models near criticality depends strongly on the system size *L*, a phenomenon known as finite-size scaling (*42*). We identified as a key experimental observable the ratio 〈Δ*t*〉 / 〈*t*〉 between the residence and transition timescales (see Supplementary Text). Simulations at various values of *J* indeed revealed a strong dependence of 〈Δ*t*〉 / 〈*τ*〉 on *L* (Fig.3a), with all results collapsing onto a single curve defined by the finite-size scaling relation 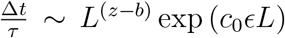 where *z, b, c*_0_ are scaling constants and *ϵ* = |*J* – *J*\*| /*J*\* is a “reduced temperature” providing a dimensionless measure of the (energetic) distance to criticality (*43, 44*) (see Supplementary Text and Fig. S9 for determination of the scaling constants). Because different combinations of *J* and *L* can yield the same value of *ϵL*, this scaling does not allow unique determination of *J* from the measured 〈Δ*t*〉 / 〈*t*〉. However, finite-size scaling theory predicts that the critical coupling energy *J*\*, separating the highly ordered (polarized) and disordered (non-polarized) regimes, also depends on the system size *L* (*45*). Consistent with this prediction, a phase diagram on the *J-L* plane constructed from simulations showed that the boundary between polarized (Fig. 3b, gray region) and non-polarized (Fig. 3b, blue region) dynamics coincided well with the theoretically predicted scaling (Fig. 3b, gold curve) 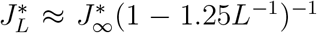, where 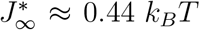 is the critical coupling energy in the thermodynamic limit (*L* → ∞) (see Supplementary Text). Interestingly, we found that iso-lines corresponding to *L-J* combinations of constant 〈Δ*t*〉 / 〈*τ*〉 (Fig.3b, beaded curves) were nearly parallel with the profile of 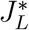. Dividing out 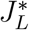 from the iso-lines corresponding to the measured 〈Δ*t*〉 / 〈*τ*〉 values of Tar (〈Δ*t*〉 / 〈*τ*〉 ≈ 8; blue in Fig. 3b) and Tsr (〈Δ*t*〉 / 〈*τ*〉 ≈ 12; red in Fig. 3b) revealed remarkable confinement of the coupling energy for both Tar and Tsr to within ±3% of 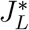 across a broad range in *L* (Fig 3b, Inset). Thus, despite uncertainty in the array size *L*, finite-size scaling analysis of the observed temporal statistics strongly suggests that both Tar and Tsr arrays are tuned very close to the Ising critical point.

**Figure 3:**
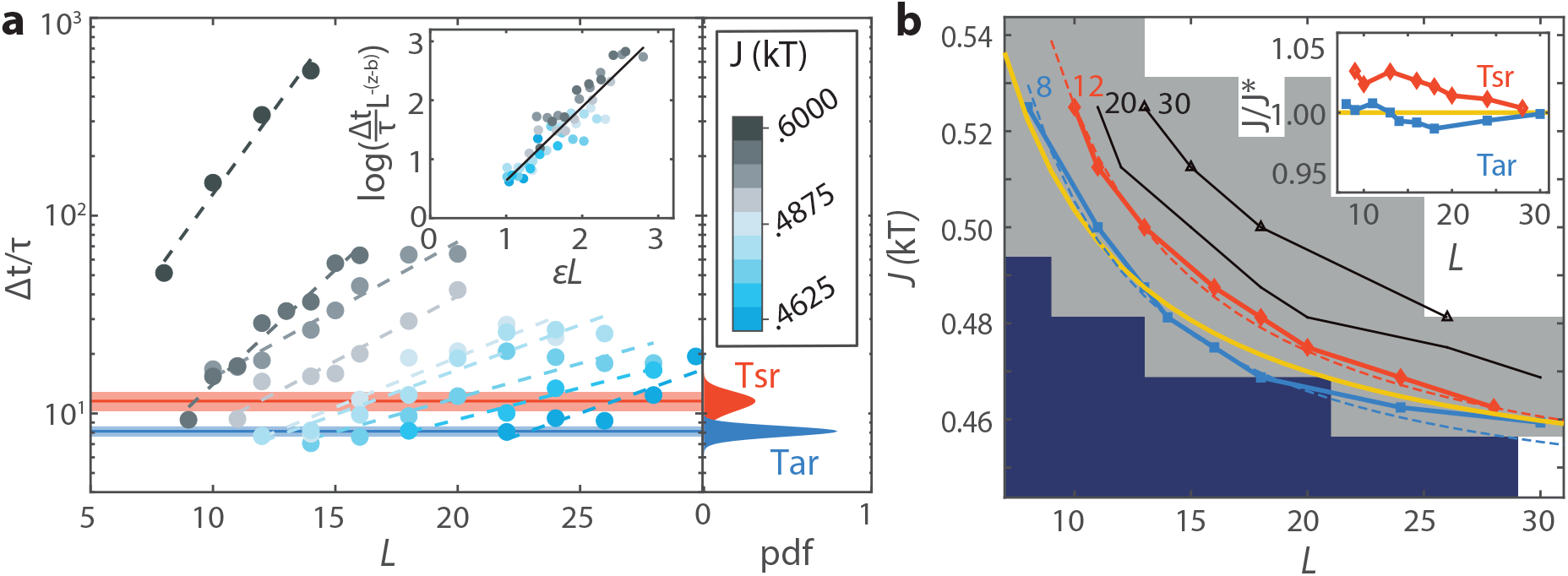
Finite-size scaling analysis of switching statistics reveals near-critical tuning of chemosensory arrays. **(a)** (left) Switching timescale ratio 〈Δ*t*〉 / 〈*τ*〉 for various values of the coupling energy *J* as a function of lattice size *L* (circles), with exponential fits (dashed lines). Inset shows data collapse for the near-critical region (*J*<0.6 *k_B_T*), upon rescaling by factors *ϵ* = | *J* – *J* * | /*J* * and *L*^−(*z*–*b*)^ according to finite-size scaling theory (see Fig. S9 for determination of scaling parameters). (right) Estimated distribution (see Fig. S10) of the switching timescale ratio 〈Δ*t*〉 / 〈*τ*〉 across cells, for Tsr-I214K (red) or Tar [QEEE] (blue). Inset: Switching timescale ratio iso-lines for Tar (blue) and Tsr, normalized by the finite size scaling of the critical coupling energy *J*\* reveals tuning within 3% of the critical energy. **(b)** Phase-diagram of switching behavior as in the *L-J* plane. Parameter value pairs generated polarized (grey region) and non-polarized (dark blue region) fluctuations as estimated from the peak-valley ratio of the time series histograms (white: not determined). Iso-lines of similar switching timescale ratios were constructed from the simulations (beaded curves). Theoretical finite-size scaling predictions for the critical coupling energy 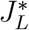 (*45*) (gold curve) and the switching time ratios (dashed lines, see Table S11 for fit parameters) are superimposed.

Finite-size scaling analysis also allowed us to estimate from the observed residence and transition times the fundamental flipping timescale 1/*ω*_0_ of allosteric units as a function of *L* (see Supplementary Text), and this depenence was very similar for both Tar and Tsr (Fig. 4b). Although our FRET measurements do not provide a direct estimate of *L*, we can motivate approximate upper and lower bounds based on structural and biochemical findings in the literature. Cryo-EM studies have revealed the detailed ultrastructure of bacterial chemosensory arrays (*24, 25, 46*), revealing an extended regular arrangement (Fig. 4a) of ‘core units’, the smallest complex of array components that has shown kinase activity *in vitro* (*47*) and *in vivo* (*48*), consisting of one CheA kinase dimer, two CheW scaffolding proteins and six trimers of chemoreceptor dimers. Expression data under defined growth conditions (*35*) suggest an approximate number of ≈ 1000 core units per cell, close in number to a lattice of size *L* × *L* =30 × 30, which we take as an approximate upper bound. However, it is also possible that the fundamental unit of cooperativity is an even larger complex than the core units. For example, taking instead the repeating unit cell of the array structure’s p6 symmetry group (Fig. 4a) as the allosteric unit leads to ≈ 300 repeating units per cell, or approximately *L* × *L* =17× 17, which we take as an approximate lower bound. With these approximate limits, our scaling analysis yields a flipping timescale of individual allosteric units in the range 1/*ω*_0_ ≈ 15-35 ms (Fig. 4b). The importance of protein structural dynamics for function is increasingly recognized (*49*), but estimates of the transition timescales are usually obtained via *in vitro* measurements (*50–52*) or molecular dynamics simulations (*53*), and span an enormously wide range (from picoseconds to milliseconds). Our *in vivo* estimate for the chemoreceptor array allosteric unit lies near the upper extreme of that range, perhaps reflecting the large size of allosteric units (even the core unit, the smaller of the two limits considered here, contains 16 protein monomers).

**Figure 4:**
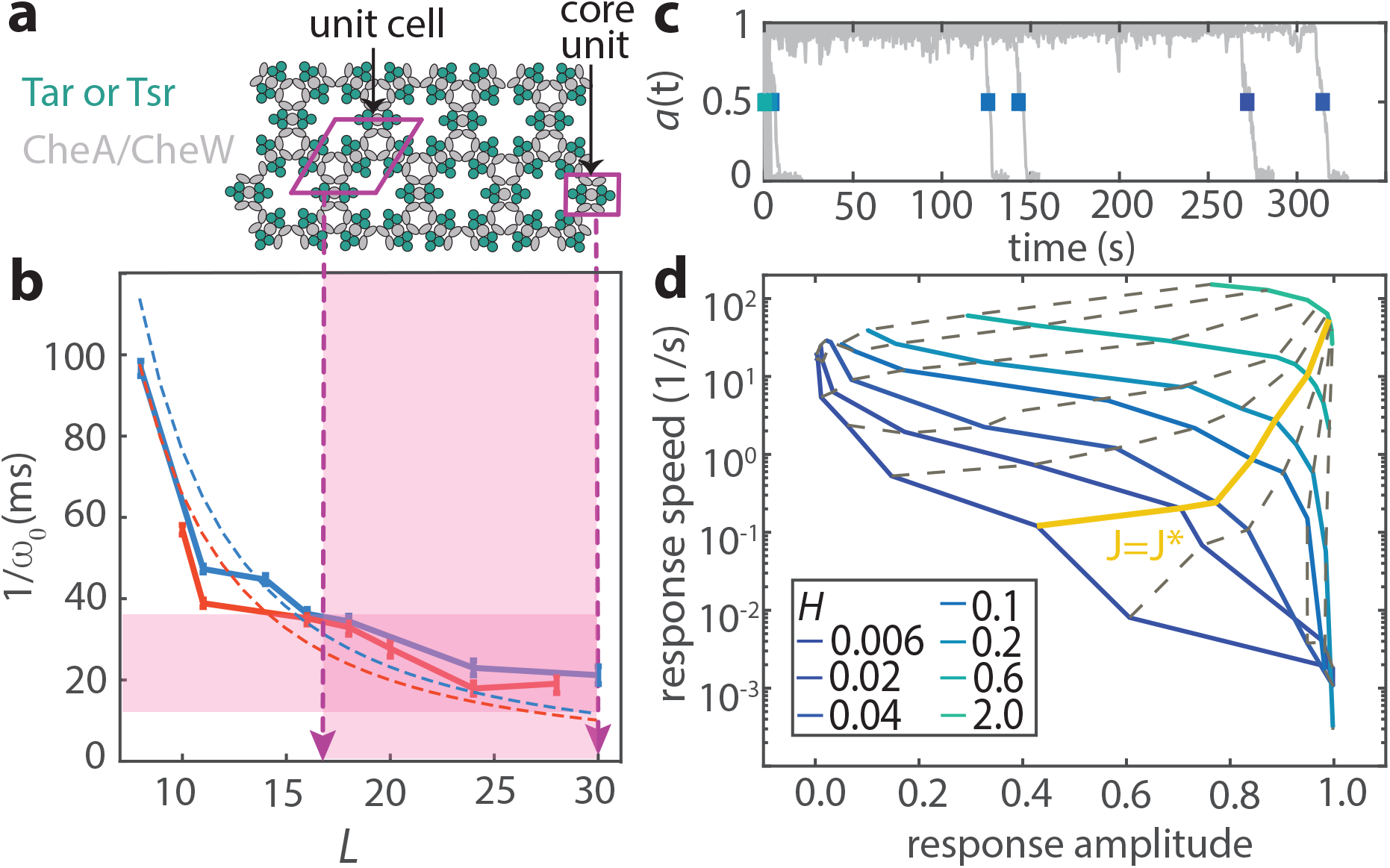
Mechanistic and functional implications of near-critical tuning: allosteric timescale and the speed-amplitude tradeoff. **(a)** Schematic top view of chemosensory array with possible fundamental units of cooperativity in the bacterial chemosensory array: core unit consisting of a CheA kinase dimer, two CheW monomers, and six trimers of chemoreceptor dimers (purple rectangle) and the repeating unit cell of the array, consisting of one full core unit and four half core units (purple parallelogram). Each chemosensory array contains approximately 1000 core units or 300 repeating unit cells. **(b)** Fundamental transition timescale 1/*ω*_0_ of allosteric units, plotted as a function of the assumed array size, *L*, for Tar (blue) or Tsr (red) arrays, based on simulations calibrated by experimentally observed residence- and transition-times (solid lines), shown together with the expected dependence based on finite sized scaling theory (dashed lines; see Supplementary Text). The lower and upper limits of array size estimates from biochemical and biophysical array characterisation (see main text) are superimposed and yield a timescale range 1/*ω*_0_ ≈15-35 ms (shaded area). **(c)** Simulated activity time series for a 20×20 array with *J* = 0.5938, varying values of *H*, all initialized with *a*=1 and assuming 1/*ω*_0_ = 25 ms (panel b). For each of the 10 time series the response time, defined as the time of the first crossing of *a*=0.5 is indicated (colored rectangle). If no crossing is detected, the maximum simulated time is used as a lower bound of the transition time. **(d)** Phase diagram showing the response amplitude 1 – 〈*a*〉 (*J, H*)/0.5 versus the array response speed, as determined from the first time for the lattice activity, initialized at *a*=1, to cross *a*=0.5 (Fig. S11). Iso-lines of constant coupling energy *J* (dashed lines and solid gold) indicate that near-critical tuning strikes a balance between response speed and amplitude for a range of external fields (colored lines).

Why are chemosensory arrays tuned so close to criticality? Previous theoretical studies have shown that signal amplification due to coupling within Ising-type conformational spread models comes at the cost of prolonged response times (*54, 55*). To address the influence of receptor coupling on response times to external stimuli, we simulated the dynamic response of arrays with *L* = 20 after an applied step stimulus (implemented within the model by an external ligand field *H_L_* that modifies the flipping rate by a factor *e^H^L* (1–2*a*); see Materials and Methods) favoring the inactive conformation to mimic the effect of attractant chemoeffector stimulation, and repeatedly (*N*=10) measured the response time (Fig. 4c, colored rectangles) as well as the response amplitude for various combinations of *J* and *H_L_* (Fig. S11). For every stimulus size *H_L_*, increasing the coupling strength *J* led to a decrease in the response speed (defined as the inverse of response time) but an increase in response amplitude, indicating a tradeoff (Fig. 4d). The profile of these *H_L_*-isolines are interesting when viewed as a Pareto front (*56*) for navigating the tradeoff, having a convex shape with a knee above which response speed drops off sharply. Remarkably, the critical coupling energy (*J* = *J*\*; Fig. 4d, yellow curve) traverses this knee at every stimulus size, indicating that near-critical tuning allows balancing of these two response objectives, allowing for large response amplitudes without drastically compromising response speed.

A compelling open question is how *E.coli* chemoreceptor arrays achieve the observed near-critical tuning. The strength of conformational interactions in canonical allosteric oligomers such as hemoglobin are usually assumed to be determined by their tertiary structure, which is in turn encoded by their amino acid sequence. While it is possible that the conformational coupling strength *J* is similarly ”hard-coded” in bacterial chemoreceptor arrays, recent experiments combining FRET with *in vivo* cross-linking revealed that the expression level of the scaffolding protein CheW can affect both the array composition and apparent response cooperativity at the population level (*57*). Future single-cell FRET experiments that monitor switching fluctuations of arrays with similarly altered compositions could reveal whether and to what extent conformational coupling within this assembly can be tuned by physiological mechanisms such as gene expression.

An increasing number of biological systems involving many components, from protein sequences (*58*) to cellular membranes (*59*), as well as communication between cells (*60, 61*) and even whole organisms (*62, 63*), have been reported to self-organize into narrow zones of phase space close to a critical point, on the boundary between order and disorder (*64*). Our finding that bacterial chemoreceptor arrays are tuned close to the Ising critical point reveals near-critical tuning as a design principle for balancing competing response objectives in large allosteric signaling assemblies. Given the many spatially extended arrays being discovered across cell biology (*4, 5, 18, 19*) as well as their anticipated role in synthetic biology (*7*), we envisage that our approach of exploiting *in vivo* noise signatures of such assemblies to understand their function could find use across a wide array of systems.

## Acknowledgments

We thank Sander Tans, Bela Mulder, Pieter Rein ten Wolde, and Kristina Ganzinger for critical reading of the manuscript. We also thank Simone Boskamp for strain construction, Mathijs Rozemuller for help with numerical simulations, AMOLF support departments for providing microscopy and electronic assistance. This work is dedicated to the memory of Howard C. Berg (1934-2021) and Thomas A. J. Duke (1964-2012).

## Author contributions

JMK and TSS conceived study. FA and JMK performed experiments. JMK and FA performed numerical simulations, with code developed by YM. JMK,FA,JSP, and TSS analyzed and interpreted data. JSP provided the chemoreceptor mutants. JMK, FA and TSS wrote the paper with input from JSP and YM.

## Supplementary materials

Materials and Methods

Supplementary Text

Figs. S1 to S11

Tables S1 to S15

References (*65–80*)

