## Supplementary material for "Near-critical tuning of cooperativity revealed by spontaneous switching in a protein signalling array": Materials & Methods; Supplementary Text, Tables and Figures

**This supplement contains:**
Materials and Methods
Supplementary Text
Tables S1 to S15
Figures S1 to S11
References *(65-80)*

### 1 Materials and Methods

#### 1.1 Cell culture and Growth Media

All experiments were performed with TSS1964 (*26*), a receptorless non-adapting derivative of *E. coli* K-12 RP437 (HCB33) expressing adhesive FliC flagellar elements that immobilizes them on glass surfaces. Chemoreceptor mutants were expressed and induced as listed in Table S1. CheZ-YFP and CheY-mRFP were expressed in tandem from pSJAB106 (*26*) using 50 μM IPTG for Tar [QEEE] and Tsr-F396Y, and 100 μM IPTG for Tsr-I214K. For experiments were CheZ localization was used to visualise chemoreceptor clusters, IPTG induction was lowered to 30 μM for Tsr-I214K. For experiments, cells were grown in Tryptone Broth ('TB':10 g/L bacto-tryptone, 5g/L NaCl) at 33.5 °C with shaking, from a saturated overnight culture in TB, with 100 μg/mL ampicillin and 34 μg/ml chloramphenicol in both cultures and appropriate inducers in the day culture. Cells were harvested at optical density 0.46 - 0.47, then washed once with and thereafter placed in motility media ('MotM': 10 mM $KPO_4$ at pH 7.0, 0.1 mM EDTA, 1 μM L-methionine, 10 mM lactic acid, 0.067 mM NaCl). Because of auxotrophic limitation of *E. coli* RP437, growth and protein expression are arrested in MotM (*65*). Cells were incubated at room temperature for 1.5 hours, before imaging, to allow for fluorescent protein maturation.

#### 1.2 FRET Microscopy

Single-cell FRET microscopy was performed essentially as reported previously (*26*). Cells were immobilized on a glass coverslip were placed in a flow cell under continuous flow (400 μL/min) established by a syringe pump (Harvard Apparatus PHD 2000, USA). The sample was illuminated every second with an LED system (CoolLED pE-2, UK) through a 40x 1.30 NA oil objective (Nikon, JP) for 17 ms. Epifluorescent light was sent through an optical splitting device (Cairn Research OptoSplit II, UK) in combination with a dichroic mirror and emission filters to project the emitted donor and emission channels in parallel on the sensor of an EM-CCD camera (Princeton Instruments proEM 512, USA), set with a multiplication gain of 100.

The flow cell temperature was controlled through a custom-built stage heater, based on a water-cooled thermoelectric Peltier element controlled through a PID controller (T = 18 °C for I214K, T = 25 °C for Tar [QEEE]). The activity bias depended on temperature (*66*), therefore we controlled the temperature to keep the average activity bias close to 1/2, at which we found the number of switchers to be maximum (Fig. S1d). The buffer solution temperature was controlled through a heat bath with heating and cooling capacity (Grant Instruments LT ecocool 100, UK).

After drift correction using rigid stack registration (*67*) in ImageJ, Images were segmented using the Mahotas python library (*26, 68*) to obtain single-cell fluorescence intensity time series. Afterwards, intensity time series were corrected for bleaching. The FRET ratio was calculated from the fluorescence time series as described previously (*20, 26*) and is proportional to kinase activity. Each ratiometric FRET time series was set to zero using the response to a saturating attractant stimulus and normalized to the maximum FRET level during a large repellent response to obtain the kinase activity within the chemosensory array per cell.

#### 1.3 Switching analysis

Switching events were analyzed automatically using a custom-made Matlab script (Mathworks, USA). For each cell, attractant and repellent responses were detected automatically based on the timing and duration of the stimulus delivery. Next, the ratiometric FRET time series of each cell, after correcting for photobleaching, was normalized between one and zero based on the repellent and attractant response amplitude, respectively. The FRET signal was then low-pass filtered using a 3s moving average filter, and switching events were detected as peaks in the derivative of the filtered signal. With each switching event, an amplitude (change in kinase activity), residence time (time until next event), and transition time (duration of the switch based on a fit of the form $1 - e^{-1}$) are associated. The switching behavior of each cell was classified according to the amplitude and number of the switching events per cell. A two-state switching cell is defined as a cell with at least $65\%$ of its transitions showing activity level changes of at least 0.7 ($70\%$ of total kinase activity). For each two-state switching cell, the bleaching correction is refined by another correction step using only the activity in either the $a = 0$ state, for cells with bias $< 0.5$, or $a = 1$ state, for cells with bias $> 0.5$. The time series is then renormalized using the maximum activity level of the histograms during the two-state switching, and the amplitude, transition time, and residence time associated with each switching event are extracted again. To verify that the automated switching analysis extracts events reliably, we generated mock time series that recapitulated the switching phenotype of Tsr-I214K and added Gaussian white noise to approximate the experimental signal-to-noise ratio. By comparing the transition and residence times of the mock time series to the times extracted by our automated analysis, we find that the relative uncertainty is minimal, even for low signal-to-noise ratios (fig.

S4).

#### 1.4 Numerical simulations of the conformational spread model

Conformational spread was modeled by a two-dimensional Ising model on an $L \times L$ lattice with free boundary conditions. Each lattice site represents an allosteric unit, whose conformational state was represented in the main text by an activity variable $a_i \in \{0, 1\}$. For the discussion here, we make the mapping $\sigma = 1 - 2a$ and discuss the model in terms of the 'spin variable' $\sigma$ which takes one of two values, $\sigma = 1$ for active, and $\sigma = -1$ for inactive. The activity of the unit at site $i$ is influenced by its $N_j$ nearest neighbours through the coupling energy $J$ a biasing field $H_b$ and a ligand field $H_L$, giving a Hamiltonian (total energy) $\mathscr{H}$ for this lattice (in units of $k_BT$):

$$\mathscr{H} = -J \sum_{\langle ij \rangle} \sigma_i \sigma_j + (H_b + H_L) \sum_i \sigma_i \tag{S1}$$

where $\langle ij \rangle$ indicates that the summation is over all nearest neighbour pairs in the lattice. The biasing and ligand fields $H_b$ and $H_L$ were set to zero for most of our simulations, except where specified otherwise in the text. The probability of finding the lattice in a given configuration is then proportional to $e^{-\mathscr{H}}$ and the ratio of probabilities $p_i(\sigma_i)$ and $p_i(-\sigma_i)$ for the $i$-th site to be in state $\sigma_i$ as opposed to $-\sigma_i$ is

$$\frac{p_i(\sigma_i)}{p_i(-\sigma_i)} = e^{-\Delta\mathscr{H}} \tag{S2}$$

where $\Delta\mathscr{H} \equiv \mathscr{H}(\sigma_i) - \mathscr{H}(-\sigma_i)$ is the change in the Hamiltonian upon flipping $\sigma_i$. We set the rate $\omega(\sigma_i \to -\sigma_i)$ for the unit at site $i$ to flip from a state $\sigma_i$ to the opposite state $-\sigma_i$ as

$$\omega(\sigma_i \to -\sigma_i) = \omega_0 \exp\left[-J\sigma_i \sum_j^{N_j} \sigma_j + (H_b + H_L)\sigma_i\right] \tag{S3}$$

where $\omega_0$ is the fundamental flipping frequency of a single lattice unit and $N_j$ is the number of nearest neighbors, so as to satisfy (together with Eqs. S1 and S2) the detailed balancing condition,

$$\frac{p_i(\sigma_i)}{p_i(-\sigma_i)} = \frac{\omega(-\sigma_i \to \sigma_i)}{\omega(\sigma_i \to -\sigma_i)}. \tag{S4}$$

We note that the fundamental frequency $\omega_0$ corresponds to the rate of conformational transitions in an individual allosteric unit. Because this rate is unknown for the bacterial chemosensory array, in our simulations we set this parameter to unity, implying that the simulated temporal statistics are expressed in units of the fundamental timescale $1/\omega_0$.

The rates as defined in Eq. S3 were used in a kinetic Monte Carlo scheme (essentially as in (*69*), but modified to have free boundary conditions; see below) which uses in every iteration one random number to draw the time until the next flip, and another to determine which site flips. The lattice was an $L \times L$ lattice with free boundary conditions, such that the number of nearest neighbors $N_j$ at each lattice site was

$$N_j = \begin{cases} 2 & \text{at corners,} \\ 3 & \text{at edges,} \\ 4 & \text{otherwise.} \end{cases} \tag{S5}$$

To calculate the activity $a$ of the lattice at each time point, we map the spin variable $\sigma_i \in \{-1, +1\}$ at each site to an activity variable $a_i \in \{0, 1\}$ as $a_i = (\sigma + 1)/2$ and take its mean across the lattice,

$$a = \frac{1}{L^2} \sum_i a_i. \tag{S6}$$

Simulations were implemented with Python (code will be made available online) and single simulation runs for extended times were performed on regular desktop computers. Parallel computations were performed on the LISA cluster of the SURFsara national computing facility (Amsterdam). For the parallel runs, each parameter set was given a unique seed for random number generation. Resulting time series were sampled at regular intervals, and subsequently downsampled to approximate the acquisition frequency of experiments (1Hz) relative to the array-level switching frequency ($\sim 10^{-2}$ Hz). Simulated time activity time series so obtained were then further processed in the same way as experimental data to extract the temporal statistics of switching.

### 2 Supplementary Text

#### 2.1 Changing activity bias by genetic modification

The activity of the chemoreceptors in bacteria is determined by fast ligand binding and slow changes in the activity bias level through modifications of the chemoreceptors by a pair of adaptation enzymes (*70*). Upon attractant binding, the activity decreases, but activity-dependent modification of the chemoreceptors provides effective negative feedback that restores the activity to its prestimulus level. This way, the wild-type chemosensory array faithfully returns to a steady-state activity bias of, on average, 0.3, but with variation between cells presumably due to expression level variation of the adaptation enzymes (*26*). In the absence of the adaptation enzymes, both chemoreceptors Tar and Tsr when natively expressed, have an activity bias close to 1 (*20*). Switching between the active and inactive array state is only observed if the activity bias has intermediate values and maximizes at 0.5. In our previous study on fluctuations in chemosensory activity (*26*), we used small amounts of chemoattractant to bring the activity bias to intermediate values to observe switching behavior. Here, we wanted to observe fluctuations in the absence of chemoeffectors and adaptation enzymes and therefore change the activity bias in a ligand-independent way.

The adaptation enzymes (CheR and CheB in *E. coli*) change the energy bias of the chemoreceptors by sequentially methylating and demethylating of 4 conserved glutamate residues on chemoreceptor Tar and 5 in Tsr (*31, 71, 72*)), where (CheR-mediated) methylation increases the activity bias and demethylation (by CheB) decreases it. Previous studies identified genetic modifications of those sites that mimic methylation. For both chemoreceptors the effect of methylation can be mimicked by exchanging a glutamate (E) for glutamine (Q). The fully unmethylated state (EEEE(E)) has an activity bias close to 0. For Tar, it QEEE modification state has an intermediate activity bias and shows fluctuations in the absence of chemoeffectors (*27*), and is used throughout this work. Unfortunately, in the case of Tsr, the QEEEE modification state has a bias close to 1 (*72*). Therefore, we pursued alternative modifications that could produce an intermediate activity bias.

Of the many available mutant Tsr receptors, Tsr-I214K in the control cable region of Tsr was a promising candidate (*32*). In population-averaged FRET experiments, it showed an intermediate activity bias and similar dose-response curve parameters to Serine as Tar [QEEE] to MeAsp. Therefore, we investigated the behavior of cells expressing only Tsr-I214K with single-cell FRET and observed similar switching behavior as in the case of Tar [QEEE].

#### 2.2 Finite-size scaling effect in Ising lattices

##### 2.2.1 The Ising model in the thermodynamic limit can not explain spontaneous switching

The two-dimensional Ising model (*73, 74*) is known to demonstrate a second-order phase transition as a function of the ratio $J/k_BT$ where $J$ is the coupling energy, $T$ is the temperature, and $k_B$ is Boltzmann's constant. Increasing $J$ across a critical value $J^*$ at fixed $T$ (or equivalently, decreasing $T$ below a critical value $T^*$ at fixed $J$) leads to an abrupt ordering of the lattice for $J$ above (or $T$ below) this critical point, with the lattice becoming polarized with all spins in either the up or down states. For a two-dimensional lattice in the thermodynamic limit (where the lattice size $L \to \infty$), an exact solution of the model was obtained by Onsager (*75*), giving the precise critical value $J^*/k_B T = \ln(1+\sqrt(2))/2 = 0.440686\ldots$ (and equivalently $k_B T^*/J = 2.26918\ldots$). Near this critical point, various thermodynamic observables (such as the correlation length $\xi$ and the correlation time $\tau_c$) are known to diverge as power laws characterized by critical exponents as a function of the "reduced temperature",

$$\epsilon \equiv |T - T^*|/T^* = |J^* - J|/J, \tag{S7}$$

which provides a dimensionless measure of the distance from the critical point (*76*). For example, the critical exponent $\nu$ ($= 1$ for 2-D lattices) defines how the correlation length diverges, $\xi \sim \epsilon^{-\nu}$, and another exponent $z$ ($\approx 2.2$ for 2-D lattices (*44*)) in turn defines the scaling of the correlation time $\tau_c \sim \xi^z = \epsilon^{-\nu z}$, as the critical point is approached ($\epsilon \to 0$). The divergence of the correlation time at the critical point implies that above $J^*$ (or equivalently below $T^*$), the system becomes frozen in one of the two polarized states. Thus, the Ising model in the thermodynamic limit is unable to explain the switching fluctutions we have observed in chemoreceptor arrays.

###### 2.2.2 Finite-size Ising models exhibit two-state switching near criticality

By contrast to the behavior in the thermodynamic limit discussed above, in Ising lattices of finite size, the correlation length $\xi$ can not grow beyond the lattice size $L$, and the scaling of various observables near criticality becomes a function of the system size $L$, a phenomenon known as finite-size scaling (*42, 77*). For example, the singularity in the correlation time scaling $\tau_c \sim \xi^z = \epsilon^{-\nu z}$ at the critical point ($\epsilon = 0$) is lost due to the cutoff of $\xi$ at $L$ and instead becomes $\tau_c \sim L^z$. Thus, fluctuations in near-critical 2-D Ising lattices of finite size demonstrate all-or-none switching between polarized states (intuitively, because $\xi \sim L$) that become increasingly slow as the lattice size grows (intuitively, because $\tau_c \sim L^z$) — an example of the phenomenon of "critical slowing down" observed across a broad range of dynamical systems near criticality. The excellent agreement between the temporal statistics of our experimental data with those of simulated Ising lattices (Fig. 2) suggested that both Tar and Tsr chemoreceptor arrays are well described as finite-size Ising systems tuned close to criticality.

To obtain a quantitative estimate for how closely tuned chemosensory arrays are to criticality, we applied finite-size scaling to further analyze our experimental and simulated temporal statistics, as described below. The basic idea is to use the observed degree of near-critical slowing to estimate the distance to criticality, and (with appropriate calibrations) also obtain an estimate for the fundamental conformational timescale $1/\omega_0$ of allosteric units.

##### 2.2.3 Scaling of the critical energy

An important result of finite-size scaling theory is that the position of the critical point (*i.e.* the value of $J^*$, or equivalently, $T^*$) itself becomes a function of the system size $L$ (*77*). As noted above, the well-known Onsager exact solution for the infinite two-dimensional Ising lattice (*75*) gives the critical coupling energy $J^*$ (in units of $k_BT$) explicitly as

$$J^*_\infty = \frac{1}{2}\ln\left(1+\sqrt{2}\right) \approx 0.44. \tag{S8}$$

For finite lattices there is no exact analytical solution, but both analytical (*78*) and numerical (*45*) approaches to finite-size scaling theory have established that the critical coupling energy $J^*$ (or equivalently the critical temperature $T^*$) dependence on the system size $L$. The dependence $J^*_L$ of the critical coupling energy with system size is well approximated as (*45, 78*),

$$J^*_L \approx \frac{J^*_\infty}{1-cL^{-1}}, \tag{S9}$$

where $c$ is constant whose value and sign depend on the boundary conditions. For periodic boundary conditions, $c$ is negative ($c_{p.b.c.} = -0.36$) and hence $J^*_L$ approaches $J^*_\infty$ from below as $L \to \infty$. By contrast, $c$ is positive for free boundary conditions ($c_{f.b.c.} = +1.25$) and in this case $J^*_L$ approaches $J^*_\infty$ from above as $L \to \infty$. Given that chemoreceptor arrays cover only a finite area of the plasma membrane, they have open ends. In our simulations, therefore, only free boundary conditions are considered, meaning that $J^*_L > J^*_\infty$ for all values of $L$.

##### 2.2.4 Scaling of the residence and transition times

As noted above, for coupling energies close to the critical energy ($J \approx J^*$), the spatial correlation length of activity states approaches the lattice size $L$, leading to polarized all-or-none fluctuations. Thus, within this regime, the activity time series is well approximated as a random telegraph process (*79*), and hence the residence timescale $\Delta t$ (i.e. time between switching events) is expected to be proportional to the correlation time $\tau_c$ (because $\tau_c = (\langle\Delta t_{\text{up}}\rangle^{-1} + \langle\Delta t_{\text{down}}\rangle^{-1})^{-1}$ for a random telegraph process). Numerical studies have found that a good approximation for the scaling of $\tau_c$ in finite lattices in the near-critical region, $\tau_c \sim L^z \exp\left[c_0\epsilon L\right]$, with $\epsilon = |(J^*_\infty - J)/J|$ and the "dynamical critical exponent" $z = 2.2 \pm 0.1$ (*43, 44*). We confirmed that the residence times extracted from our simulations at various combinations of $\epsilon$ and $L$ were well-fit by a scaling of the same form,

$$\Delta t = c_t L^z \exp\left[c_0\epsilon L\right] \tag{S10}$$

with $z = 2.2$ (Fig. S9a), and these fits yielded for the remaining scaling constants $c_0 = 1.6 \pm 0.5$ and $c_t = 1.4 \pm 0.5$.

For the transition time $\tau$ (the duration of the activity transient upon switching), we found that our data were well-described (Fig S9b) by a power-law,

$$\tau = c_\tau L^b \tag{S11}$$

with $c_\tau = 1$ and $b = 1.725$.

##### 2.2.5 Scaling analysis of the experimental data

While the finite-size scaling relations obtained above provide an excellent approximation to the simulated data, they can not be directly applied to the experimentally observed timescales because the simulations yield timescales in units of the fundamental flipping timescale $1/\omega_0$, which remains unknown for the chemosensory array. That is, $\Delta t_{\text{sim}}$ and $\tau_{\text{sim}}$ from simulatios are related to $\Delta t_{\text{exp}}$ and $\tau_{\text{exp}}$ from experiments as, respectively,

$$\Delta t_{\text{sim}} = \Delta t_{\text{exp}} \omega_0, \quad \text{and} \quad \tau_{\text{sim}} = \tau_{\text{exp}} \omega_0. \tag{S12}$$

We therefore identify as a key experimental observable the dimensionless timescale ratio $\Delta t/\tau$, which removes the dependence on the unknown constant $\omega_0$. Using Eqs. S10 and S11, we obtain a scaling for the timescale ratio $\Delta t/\tau$,

$$\frac{\Delta t}{\tau} = \frac{c_t}{c_\tau} L^{(z-b)} \exp\left(c_0 \epsilon L\right). \tag{S13}$$

This scaling provides a one-to-one relationship between the timescale ratio $\Delta t/\tau$ and the product $\epsilon L$, as evidenced by the collapse of simulated data at various combinations of $\epsilon$ and $L$ onto a single curve (Fig. 3a Inset). However, because the size $L$ is not precisely known for the chemoreceptor array, we can not uniquely determine $\epsilon$.

We therefore chose to assess the distance from criticality by comparing iso-lines in the $J$-$L$ plane, corresponding to parameter combinations that yield a given value of the timescale ratio $r_{\text{iso}} \equiv \Delta t/\tau$, with $J_L^*$, with the finite-size scaling of the critical coupling energy given by Eq. S9. The iso-lines determined from simulations (Fig. 3b) were fit well by an expression obtained by solving Eq. S13 for $\epsilon$,

$$\epsilon = \frac{\log\left[r_{\text{iso}}/c_R\right] + (b-z)\log L}{c_0 L}. \tag{S14}$$

where we have defined $c_R \equiv c_t/c_\tau$. For the scaling shown in Fig. 3e (dashed curves), we fitted Eq. S14 to the data corresponding to the iso-line for Tar ($r_{\text{iso}}$=8.5) and Tsr ($r_{\text{iso}}$=11.5), where each of the parameters (Table S11) was constrained to the obtained scaling value +/- one standard deviation.

##### 2.2.6 Scaling of the fundamental frequency $\omega_0$

As noted above, the 'timescales' determined from the simulations are a dimensionless quantity $\Delta t_{\text{sim}}$, expressed relative to the fundamental flipping timescale $1/\omega_0$ of individual lattice units.

Thus, $\Delta t_{\text{sim}}$is related to experimentally observed timescale $\Delta t_{\text{exp}}$ as $\Delta t_{\text{sim}} = \Delta t_{\text{exp}} \omega_0$. Given the finite-size scaling relations obtained above, one could therefore estimate $\omega_0$ as a function of $L$ by comparing the timescales extracted from experiments and numerical simulations. To obtain such a scaling relation for $\omega_0$, we use the fact that along the iso-lines for $\Delta t/\tau$, $\Delta t$ scales in the same way as $\tau$. Thus plugging the iso-line scaling of $\epsilon$ (Eq. S14) into the residence time scaling (Eq. S10) yields for the simulated residence time,

$$\Delta t_{\text{sim}} = c_t L^z \exp\left[\log\left(c_\tau r_{\text{iso}}/c_t\right) + (b - z)\log L\right] = c_\tau r_{\text{iso}} L^b, \tag{S15}$$

which indeed scales as the transition time ($\sim L^b$), and using this we obtain the scaling relation for $\omega_0$ (as defined in Eq. S3) as a function of the lattice size $L$ and experimental residence timescale $\Delta t_{\text{exp}}$:

$$\omega_0 = \Delta t_{\text{sim}}/\Delta t_{\text{exp}} = c_\tau r_{\text{iso}} \frac{L^b}{\Delta t_{\text{exp}}}. \tag{S16}$$

### 3 Supplemental Tables

Table S1: Plasmids used in this study to express various chemoreceptor alleles.

| **Product** | **Vector** | **Induction** | **Resistance** | **Source** |
|---|---|---|---|---|
| Tar [QEEE] | pLC113 | 2.0 $\mu$M NaSal | Cam | (*71*) |
| Tar [QEEE]/R69H | pLC113 | 2.0 $\mu$M NaSal | Cam | This Study |
| Tsr-I214K | pPA114 | 0.6 $\mu$M NaSal | Cam | (*32*) |
| Tsr-I214K/R69E | pPA114 | 0.6 $\mu$M NaSal | Cam | This Study |
| Tsr-F396Y | pPA114 | 0.6 $\mu$M NaSal | Cam | (*80*) |

Table S2: Mean residence times and number of events extracted from cells expressing Tsr-I214K.

| Bias | $\langle \Delta t_{\mathrm{up}} \rangle$ (s) | $\pm$ std | $\langle \Delta t_{\mathrm{down}} \rangle$ (s) | $\pm$ std | $N_{\mathrm{up}}$ | $N_{\mathrm{down}}$ |
|---|---|---|---|---|---|---|
| $0.65 < \alpha \leq 1$ | $116.2 \pm 3.27$ | $\pm$ 101.8 | $38.6 \pm 0.79$ | $\pm$ 26.2 | 967 | 1091 |
| $0.55 < \alpha \leq 0.65$ | $72.4 \pm 2.31$ | $\pm$ 61.2 | $48.0 \pm 1.20$ | $\pm$ 32.6 | 703 | 733 |
| $0.45 < \alpha \leq 0.55$ | $60.4 \pm 1.84$ | $\pm$ 50.9 | $59.6 \pm 1.68$ | $\pm$ 46.6 | 766 | 767 |
| $0.35 < \alpha \leq 0.45$ | $56.7 \pm 2.36$ | $\pm$ 47.2 | $80.7 \pm 3.48$ | $\pm$ 68.2 | 398 | 384 |
| $0 \leq \alpha \leq 0.35$ | $41.3 \pm 1.65$ | $\pm$ 27.9 | $118.9 \pm 6.56$ | $\pm$ 104.8 | 285 | 255 |
| $0 \leq \alpha \leq 1$ | $78.2 \pm 1.36$ | $\pm$ 75.9 | $57.0 \pm 0.95$ | $\pm$ 53.9 | 3119 | 3230 |

Table S3: Mean transition times and number of events extracted from cells expressing Tsr-I214K.

| Bias | $\langle \tau_{\mathrm{up}} \rangle$ (s) | $\pm$ std | $\langle \tau_{\mathrm{down}} \rangle$ (s) | $\pm$ std | $N_{\mathrm{up}}$ | $N_{\mathrm{down}}$ |
|---|---|---|---|---|---|---|
| $0.65 < \alpha \leq 1$ | $4.28 \pm 0.10$ | $\pm$ 3.16 | $6.09 \pm 0.12$ | $\pm$ 3.49 | 970 | 837 |
| $0.55 < \alpha \leq 0.65$ | $4.31 \pm 0.12$ | $\pm$ 3.21 | $6.07 \pm 0.15$ | $\pm$ 3.57 | 660 | 572 |
| $0.45 < \alpha \leq 0.55$ | $4.18 \pm 0.12$ | $\pm$ 3.02 | $5.89 \pm 0.14$ | $\pm$ 3.54 | 681 | 604 |
| $0.35 < \alpha \leq 0.45$ | $4.36 \pm 0.17$ | $\pm$ 3.16 | $6.18 \pm 0.20$ | $\pm$ 3.60 | 362 | 320 |
| $0 \leq \alpha \leq 0.35$ | $4.49 \pm 0.20$ | $\pm$ 3.24 | $6.35 \pm 0.22$ | $\pm$ 3.32 | 262 | 235 |
| $0 \leq \alpha \leq 1$ | $4.29 \pm 0.06$ | $\pm$ 3.15 | $6.07 \pm 0.07$ | $\pm$ 3.52 | 2935 | 2568 |

Table S4: Mean residence times and number of events extracted from cells expressing Tar [QEEE].

| Bias | $\langle \Delta t_{\text{up}} \rangle$ (s) | $\pm$ std | $\langle \Delta t_{\text{down}} \rangle$ (s) | $\pm$ std | $N_{\text{up}}$ | $N_{\text{down}}$ |
|---|---|---|---|---|---|---|
| $0.65 < \alpha \leq 1$ | 79.4 ± 2.87 | ± 70.6 | 28.2 ± 0.58 | ± 14.8 | 607 | 647 |
| $0.55 < \alpha \leq 0.65$ | 56.5 ± 2.15 | ± 41.5 | 37.4 ± 1.22 | ± 23.7 | 373 | 379 |
| $0.45 < \alpha \leq 0.55$ | 44.7 ± 1.66 | ± 30.7 | 45.3 ± 1.68 | ± 31.1 | 343 | 341 |
| $0.35 < \alpha \leq 0.45$ | 39.0 ± 2.06 | ± 29.6 | 58.0 ± 3.56 | ± 50.1 | 207 | 198 |
| $0 \leq \alpha \leq 0.35$ | 32.0 ± 1.33 | ± 18.0 | 86.0 ± 5.94 | ± 77.0 | 183 | 168 |
| $0 \leq \alpha \leq 1$ | 57.5 ± 3.03 | ± 52.7 | 42.6 ± 1.95 | ± 39.3 | 1713 | 1733 |

Table S5: Mean transition times and number of events extracted from cells expressing Tar [QEEE].

| Bias | $\langle \tau_{\text{up}} \rangle$ (s) | $\pm$ std | $\langle \tau_{\text{down}} \rangle$ (s) | $\pm$ std | $N_{\text{up}}$ | $N_{\text{down}}$ |
|---|---|---|---|---|---|---|
| $0.65 < \alpha \leq 1$ | 4.70 ± 0.14 | ± 3.22 | 5.73 ± 0.15 | ± 3.33 | 556 | 501 |
| $0.55 < \alpha \leq 0.65$ | 5.13 ± 0.18 | ± 3.39 | 6.00 ± 0.21 | ± 3.42 | 346 | 271 |
| $0.45 < \alpha \leq 0.55$ | 4.71 ± 0.20 | ± 3.50 | 6.39 ± 0.20 | ± 3.18 | 301 | 260 |
| $0.35 < \alpha \leq 0.45$ | 4.81 ± 0.27 | ± 3.53 | 6.26 ± 0.28 | ± 3.35 | 176 | 140 |
| $0 \leq \alpha \leq 0.35$ | 4.56 ± 0.24 | ± 3.09 | 6.61 ± 0.31 | ± 3.49 | 171 | 130 |
| $0 \leq \alpha \leq 1$ | 4.79 ± 0.08 | ± 3.34 | 6.06 ± 0.09 | ± 3.35 | 1550 | 1302 |

Table S6: Fit parameters of mean residence times per cell as a function of energy bias ΔG.

| | slope $\gamma_{\text{down}}$ | slope $\gamma_{\text{up}}$ | crossover point $\tau_{\text{up}} = \tau_{\text{down}}$ | $N$ |
|---|---|---|---|---|
| Tar [QEEE] | $-0.44$ | 0.45 | 47.0 ± 1 s | 204 |
| Tsr-I214K | $-0.39$ | 0.45 | 65.5 ± 1 s | 549 |

Table S7: Mean residence times and number of events extracted from numerical simulations using a conformational spread model with a lattice size of 20 × 20 spins and coupling energy J = 0.475 $k_B T$.

| External Field | Bias | $\langle \Delta t_{\text{up}} \times \omega \rangle$ | $\pm$ std | $\langle \Delta t_{\text{down}} \times \omega \rangle$ | $\pm$ std | $N_{\text{up}}$ | $N_{\text{down}}$ |
|---|---|---|---|---|---|---|---|
| H = $-0.006$ | $\alpha \approx 0.79$ | 4790 ± 381.43 | ± 4110 | 1250 ± 76.81 | ± 840 | 116 | 121 |
| H = $-0.002$ | $\alpha \approx 0.63$ | 2820 ± 172.44 | ± 2220 | 1680 ± 95.29 | ± 1240 | 166 | 168 |
| H = 0 | $\alpha \approx 0.51$ | 2190 ± 128.34 | ± 1600 | 2130 ± 149.60 | ± 1870 | 155 | 156 |
| H = 0.002 | $\alpha \approx 0.44$ | 1930 ± 97.63 | ± 1280 | 2440 ± 155.67 | ± 2040 | 172 | 171 |
| H = 0.006 | $\alpha \approx 0.23$ | 1370 ± 82.68 | ± 920 | 4480 ± 359.98 | ± 3890 | 123 | 117 |
| | | 2185 ± 90.52 | ± 1597 | 2132 ± 87.50 | ± 1868 | 732 | 733 |

Table S8: Mean transition times and number of events extracted from numerical simulations using a conformational spread model with a lattice size of 20 $\times$ 20 spins and coupling energy J = 0.475 $k_B T$.

| External Field | Bias | $\langle\tau_{\text{up}}\times\omega\rangle$ | $\pm$ std | $\langle\tau_{\text{down}}\times\omega\rangle$ | $\pm$ std | $N_{\text{up}}$ | $N_{\text{down}}$ |
|---|---|---|---|---|---|---|---|
| H = $-0.006$ | $\alpha \approx 0.79$ | 179.76 $\pm$ 10.35 | $\pm$ 111.51 | 169.54 $\pm$ 11.05 | $\pm$ 118.01 | 116 | 114 |
| H = $-0.002$ | $\alpha \approx 0.63$ | 169.18 $\pm$ 8.83 | $\pm$ 113.11 | 166.95 $\pm$ 9.16 | $\pm$ 115.50 | 164 | 159 |
| H = 0 | $\alpha \approx 0.51$ | 166.57 $\pm$ 8.95 | $\pm$ 106.98 | 156.20 $\pm$ 9.24 | $\pm$ 113.49 | 143 | 151 |
| H = 0.002 | $\alpha \approx 0.44$ | 184.13 $\pm$ 9.74 | $\pm$ 124.39 | 155.91 $\pm$ 7.38 | $\pm$ 95.05 | 163 | 166 |
| H = 0.006 | $\alpha \approx 0.23$ | 154.32 $\pm$ 8.79 | $\pm$ 91.73 | 159.81 $\pm$ 10.54 | $\pm$ 113.00 | 109 | 115 |
| | | 171.59 $\pm$ 8.79 | $\pm$ 111.49 | 161.30 $\pm$ 10.54 | $\pm$ 110.41 | 695 | 705 |

Table S9: Mean residence times and number of events extracted from numerical simulations using a conformational spread model with a lattice size of 12 $\times$ 12 spins and coupling energy J = 0.5 $k_B T$.

| External Field | Bias | $\langle\Delta t_{\text{up}}\times\omega\rangle$ | $\pm$ std | $\langle\Delta t_{\text{down}}\times\omega\rangle$ | $\pm$ std | $N_{\text{up}}$ | $N_{\text{down}}$ |
|---|---|---|---|---|---|---|---|
| H = $-0.02$ | $\alpha \approx 0.85$ | 2230 $\pm$ 96.18 | $\pm$ 2120 | 390 $\pm$ 11.34 | $\pm$ 250 | 486 | 491 |
| H = $-0.006$ | $\alpha \approx 0.66$ | 1040 $\pm$ 45.69 | $\pm$ 930 | 520 $\pm$ 21.05 | $\pm$ 430 | 412 | 410 |
| H = 0 | $\alpha \approx 0.53$ | 800 $\pm$ 31.53 | $\pm$ 650 | 700 $\pm$ 31.20 | $\pm$ 640 | 419 | 420 |
| H = 0.006 | $\alpha \approx 0.40$ | 610 $\pm$ 24.81 | $\pm$ 510 | 920 $\pm$ 38.20 | $\pm$ 780 | 416 | 415 |
| H = 0.02 | $\alpha \approx 0.22$ | 520 $\pm$ 27.07 | $\pm$ 640 | 1810 $\pm$ 69.06 | $\pm$ 1610 | 551 | 546 |
| | | 904 $\pm$ 27.60 | $\pm$ 1318 | 1045 $\pm$ 22.34 | $\pm$ 1067 | 2284 | 2282 |

Table S10: Mean transition times and number of events extracted from numerical simulations using a conformational spread model with a lattice size of 12 $\times$ 12 spins and coupling energy J = 0.5 $k_B T$.

| External Field | Bias | $\langle\tau_{\text{up}}\times\omega\rangle$ | $\pm$ std | $\langle\tau_{\text{down}}\times\omega\rangle$ | $\pm$ std | $N_{\text{up}}$ | $N_{\text{down}}$ |
|---|---|---|---|---|---|---|---|
| H = -0.02 | $\alpha \approx 0.85$ | 56.05 $\pm$ 1.89 | $\pm$ 40.04 | 57.59 $\pm$ 1.97 | $\pm$ 41.06 | 450 | 434 |
| H = -0.006 | $\alpha \approx 0.66$ | 58.07 $\pm$ 1.94 | $\pm$ 38.02 | 60.16 $\pm$ 2.11 | $\pm$ 41.10 | 385 | 380 |
| H = 0 | $\alpha \approx 0.53$ | 59.27 $\pm$ 2.17 | $\pm$ 43.31 | 54.42 $\pm$ 1.87 | $\pm$ 37.14 | 398 | 394 |
| H = 0.006 | $\alpha \approx 0.40$ | 58.84 $\pm$ 1.98 | $\pm$ 38.48 | 57.51 $\pm$ 2.14 | $\pm$ 41.90 | 378 | 382 |
| H = 0.02 | $\alpha \approx 0.22$ | 60.04 $\pm$ 1.99 | $\pm$ 44.15 | 54.77 $\pm$ 1.60 | $\pm$ 36.41 | 492 | 516 |
| | | 58.46 $\pm$0.89 | $\pm$ 41.03 | 56.76 $\pm$ 0.86 | $\pm$ 39.43 | 2103 | 2106 |

Table S11: Fit parameters of exponential fits to residence times extracted from cells expressing Tsr-I214K. Exponential fit of the form $\alpha e^{\beta x}$.

| Bias | $\Delta t_{up}$ | $\Delta t_{down}$ |
|---|---|---|
| $0.65 < \alpha \leq 1$ | $\alpha$ = 0.0083<br>$\beta$ = -0.0088 | $\alpha$ = 0.0460<br>$\beta$ = -0.0329 |
| $0.55 < \alpha \leq 0.65$ | $\alpha$ = 0.0177<br>$\beta$ = -0.0166 | $\alpha$ = 0.0289<br>$\beta$ = -0.0227 |
| $0.45 < \alpha \leq 0.55$ | $\alpha$ = 0.0249<br>$\beta$ = -0.0221 | $\alpha$ = 0.0216<br>$\beta$ = -0.0185 |
| $0.35 < \alpha \leq 0.45$ | $\alpha$ = 0.0310<br>$\beta$ = -0.0273 | $\alpha$ = 0.0139<br>$\beta$ = -0.0122 |
| $0 \leq \alpha \leq 0.35$ | $\alpha$ = 0.0423<br>$\beta$ = -0.0318 | $\alpha$ = 0.0073<br>$\beta$ = -0.0065 |

Table S12: Fit parameters of exponential fits to residence times extracted from cells expressing Tar [QEEE]. Exponential fit of the form $\alpha e^{\beta x}$.

| Bias | $\Delta t_{up}$ | $\Delta t_{down}$ |
|---|---|---|
| $0.65 < \alpha \leq 1$ | $\alpha$ = 0.0154<br>$\beta$ = -0.0150 | $\alpha$ = 0.0862<br>$\beta$ = -0.0504 |
| $0.55 < \alpha \leq 0.65$ | $\alpha$ = 0.0231<br>$\beta$ = -0.0199 | $\alpha$ = 0.0480<br>$\beta$ = -0.0342 |
| $0.45 < \alpha \leq 0.55$ | $\alpha$ = 0.0360<br>$\beta$ = -0.0284 | $\alpha$ = 0.0317<br>$\beta$ = -0.0243 |
| $0.35 < \alpha \leq 0.45$ | $\alpha$ = 0.0556<br>$\beta$ = -0.0403 | $\alpha$ = 0.0227<br>$\beta$ = -0.0194 |
| $0 \leq \alpha \leq 0.35$ | $\alpha$ = 0.0746<br>$\beta$ = -0.0473 | $\alpha$ = 0.0143<br>$\beta$ = -0.0142 |

Table S13: Fit parameters of exponential fits to residence times extracted from numerical simulations using a conformational spread model with a lattice size of 20 $\times$ 20 spins and coupling energy of J = 0.475 $k_B T$. Exponential fit of the form $\alpha e^{\beta x}$.

| Bias | $\Delta t_{up}$ | $\Delta t_{down}$ |
|---|---|---|
| $H$ = -0.006 | $\alpha$ = 0.2280<br>$\beta$ = -0.2339 | $\alpha$ = 3.3210<br>$\beta$ = -1.5900 |
| $H$ = -0.002 | $\alpha$ = 0.5589<br>$\beta$ = -0.4987 | $\alpha$ = 1.4930<br>$\beta$ = -1.0050 |
| $H$ = 0 | $\alpha$ = 0.7966<br>$\beta$ = -0.6830 | $\alpha$ = 1.3090<br>$\beta$ = -0.9488 |
| $H$ = 0.002 | $\alpha$ = 0.8822<br>$\beta$ = -0.6558 | $\alpha$ = 0.6276<br>$\beta$ = -0.5173 |
| $H$ = 0.006 | $\alpha$ = 1.0670<br>$\beta$ = -0.9160 | $\alpha$ = 0.1771<br>$\beta$ = -0.0278 |

Table S14: Fit parameters of exponential fits to residence times extracted from numerical simulations using a conformational spread model with a lattice size of 12 $\times$ 12 spins and coupling energy of J = 0.5 $k_B T$. Exponential fit of the form $\alpha e^{\beta x}$.

| Bias | $\Delta t_{up}$ | $\Delta t_{down}$ |
|---|---|---|
| $H$ = -0.02 | $\alpha$ = 0.5876<br>$\beta$ = -0.5844 | $\alpha$ = 3.3820<br>$\beta$ = -3.4340 |
| $H$ = -0.006 | $\alpha$ = 1.4200<br>$\beta$ = -1.2040 | $\alpha$ = 2.4260<br>$\beta$ = -2.3460 |
| $H$ = 0 | $\alpha$ = 2.1750<br>$\beta$ = -1.6820 | $\alpha$ = 2.0490<br>$\beta$ = -1.7660 |
| $H$ = 0.006 | $\alpha$ = 3.3800<br>$\beta$ = -2.3630 | $\alpha$ = 1.0920<br>$\beta$ = -1.0250 |
| $H$ = 0.02 | $\alpha$ = 4.1970<br>$\beta$ = -3.2720 | $\alpha$ = 0.8545<br>$\beta$ = -0.8116 |

Table S15: Parameters that describe the scaling of the timescale ratio with lattice size and critical energy (Eq. S14). Each parameter is determined individually from the scaling behavior of residence and transition times (Fig. S9). For the scaling relation of the phase diagram (Fig 3), the finite scaling relation was fitted to the iso lines corresponding to Tar, where each fit parameter was allowed to vary according to the fit uncertainty of the individual fits.

| Parameter | From fitting (Fig. S9) | Used in iso-line fit (Fig. 3) | Literature |
|---|---|---|---|
| $z$ | 2.2 | 2.1 | 2.2 ± 0.1 (*44*) |
| $b$ | 1.725 ± 0.032 | 1.75 | |
| $c_1$ | 0.294 ± 0.366 | | |
| $c_2 = c_0$ | 1.608 ± 0.466 | 1.18 | |
| $c_t = e^{c_1}$ | 1.4 ± 0.5 | | |
| $c_\tau$ | 1 (assumed, not fit) | | |
| $c_R = c_t/c_\tau$ | 1.03± 0.23 | 0.8 | |

### 4 Supplemental Figures

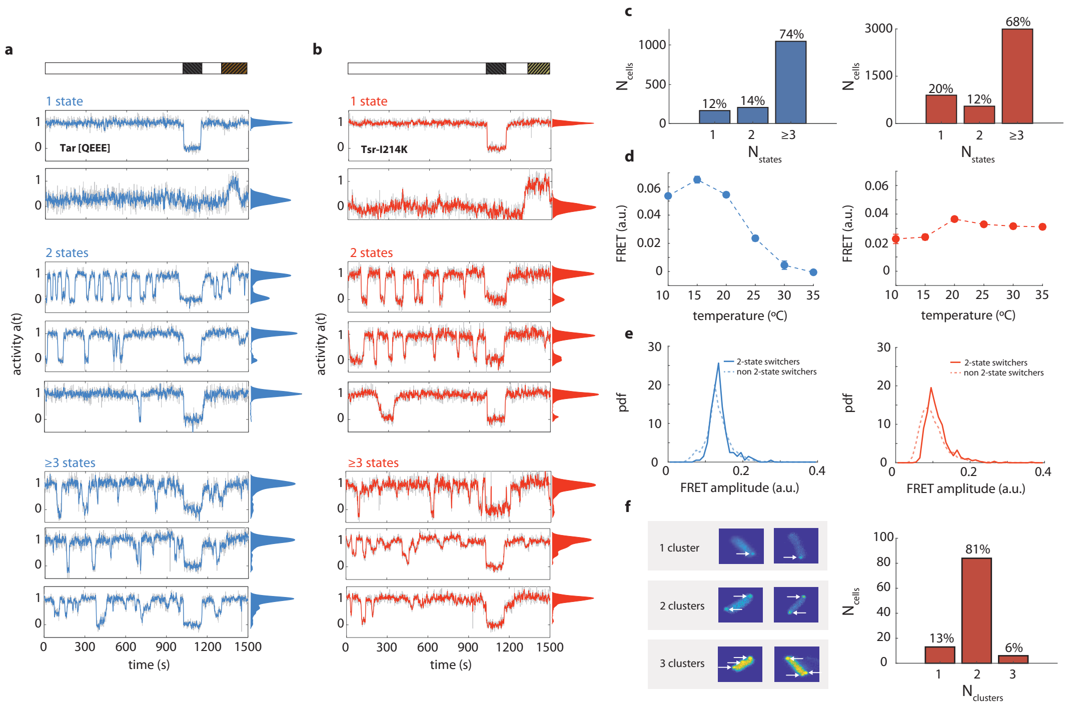


Figure **S1**: **Classification of switching behavior and temperature dependence in cells expressing chemoreceptors Tar or Tsr. (a)** (top) Stimulation protocol of the FRET experiment. Cells are exposed to measurement buffer (white) and then to an attractant stimulus of 1 mM MeAsp (dark, dashed) and a repellent stimulus of 0.3 mM $NiCl_2$ (gold, dashed). (bottom) Activity time series of eight representative individual cells expressing only the chemoreceptor Tar [QEEE] without adaptation enzymes (pVS120/TSS1964). Cells with one, two, and three or more stable states are shown. On each raw time series (grey), a 3s moving average filter was applied and the resulted time series are superimposed (blue), shown together with associated activity histograms of the filtered time series. **(b)** As in (a), but with cells expressing only Tsr-I214K in the same background (pPA114/TSS1964), and stimulated with an attractant stimulus of 1 mM Ser and a repellent stimulus of 1 mM Leu. **(c)** Histogram of number of states of total 1414 Tar [QEEE] cells (left, blue) and 4446 Tsr-I214K cells (right, red). **(d)** Temperature dependence of the steady-state FRET signal for cells expressing only Tar [QEEE] (left, blue) or Tsr-I214K (right, red). **(e)** Histrograms of maximum FRET amplitude of 1414 cells expressing Tar [QEEE] (left, blue) and 4446 cells expressing Tsr-I214K (right, red). Cells exhibiting two-state switching behavior (solid lines) have approximately equal or higher FRET responses compared to the rest of the cells (dashed lines). Maximum FRET amplitude is proportional to kinase activity. **(f)** Histogram of the number of clusters of 103 Tsr-I214K cells and example heatmaps of 6 representative cells categorized by their cluster number. FRET plasmid expression was lowered to decrease total background fluorescence and therefore allow visualization of large chemoreceptor clusters as a consequence of CheZ localization near the chemoreceptors. The percentage of cells with a single large cluster matches that of the two-state switching cells.

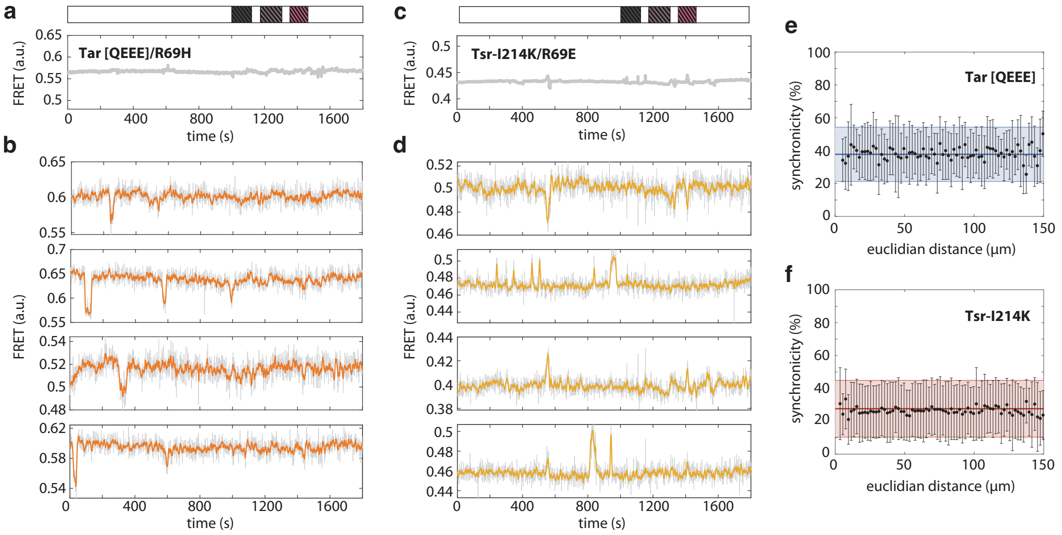


Figure **S2**: **Switching behavior does not stem from ligand sensing or cell-to-cell communication.** **(a)** Population-averaged FRET response from cells expressing only Tar Tar [QEEE]/R69H, a double mutant receptor with impaired ligand binding sites, exposed to buffer (bar, white) or different chemoeffectors (bar, shaded boxes). The following saturating doses of chemoattractants are tested here: black box, 1 mM $\alpha$-Methyl-DL-aspartic acid; grey box, 1 mM Maltose; pink box, 1 mM Leucine. Measurement at 25°C. **(b)** Single-cell time series from the experiment in (a). On each raw time series (grey), a 7s moving average filter was applied and the resulted time series are superimposed (red). **(c)** Population-averaged FRET response from cells expressing only the receptor Tsr-I214K/R69E, a double mutant receptor with impaired ligand binding sites, exposed to buffer (bar, white) or different chemoeffectors (bar, shaded boxes). The following following saturating doses of chemoattractants are tested here: black box, 1 mM L-Serine; grey box, 10 mM Cysteine; pink box, 10 mM $\alpha$-aminoisobutyric acid. Measurement at 35°C. **(d)** Single-cell time series from the experiment in (c). On each raw time series (grey), a 7s moving average filter was applied and the resulted time series are superimposed (orange). **(e)** Measure of synchronicity between 2-state switching cells expressing Tar [QEEE]. For each independent experiment, one reference cell is chosen, and the time of incidence of its switching events is compared with the time of incidence of the switching events in the rest of the cells within a window of $\pm$10 seconds. Synchronicity of 100% corresponds to all switching events coinciding within the same respective window. The process is then repeated for all cells within the same experiment and then, for all experiments. The process is repeated one more time, where switching events across all cells from the same experiment are shuffled and a number of events, corresponding to the number of events in the reference cell, are drawn randomly and with replacement. Then, the time of incidence of the reference cell's switching events is compared to this random set of events (solid blue line). Shaded area corresponds to standard deviation. The synchronicity of switching events between cells within the same experiment can be fully explained by random synchronization of switching events. **(f)** as in panel (e), but with cells expressing Tsr-I214K.

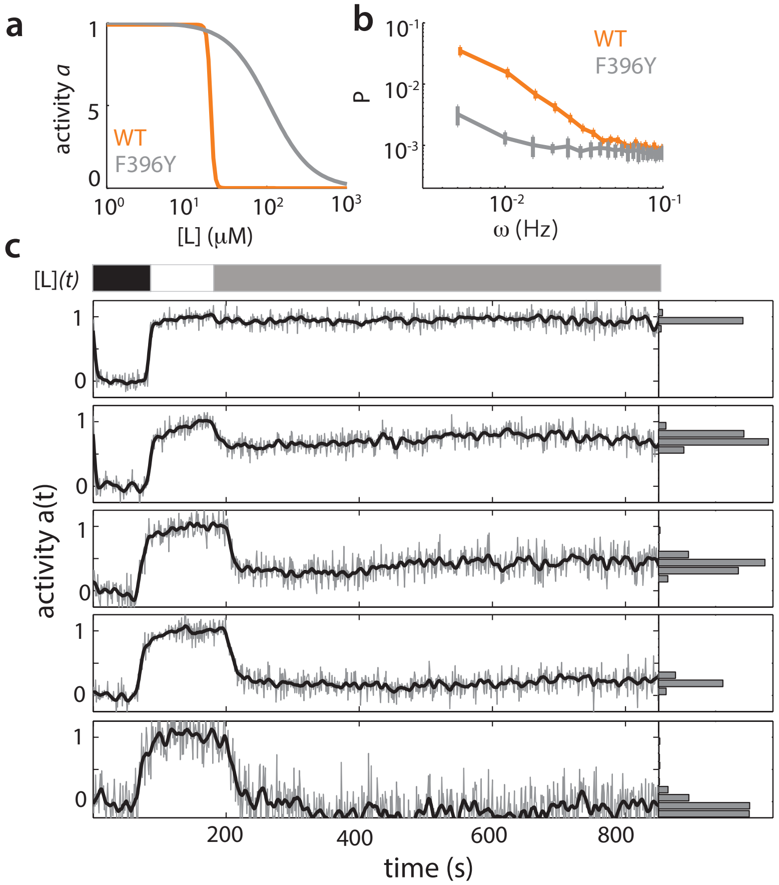


Figure **S3**: **Experiments with Tsr mutant defective in cooperativity show no stochastic switching behaviour.** **(a)** Hill curves illustrating the difference in cooperativity of the dose-response curves between wildtype Tsr and Tsr-F396Y. The dose response parameters are the average single-cell fit values obtained from single-cell FRET experiments on both genotypes in *cheRB* background. **(b)** Power specctral density (PSD) estimates for temporal signal fluctuations from cells expressing wildtype Tsr and Tsr-F396Y in cells lacking other chemoreceptors and the adaptation enzymes (TSS1964/pPA114). Shown are the average of the single-cell PSD estimates from a single experiment, error bars represent s.e.m. **(c)** (Top) Stimulus protocol for L-serine concentration ($[L](t)$). At the start of the experiment, a saturating concentration ($[L] = 1$mM, black) is applied for a short time. After flushing buffer ($[L] = 0$, white), an intermediate concentration ($[L] = 150 - 175\mu$M, gray) is sustained for $\sim$10 minutes. (Bottom) Representative single-cell time series, each normalized to its activity level before adding the first stimulus. The top two curves are taken from an experiment (34 cells) responding to 150μM, bottom three curves from a second experiment (49 cells) responding to 175μM. To the unfiltered data (gray) a 10s moving average filter is applied and superimposed.

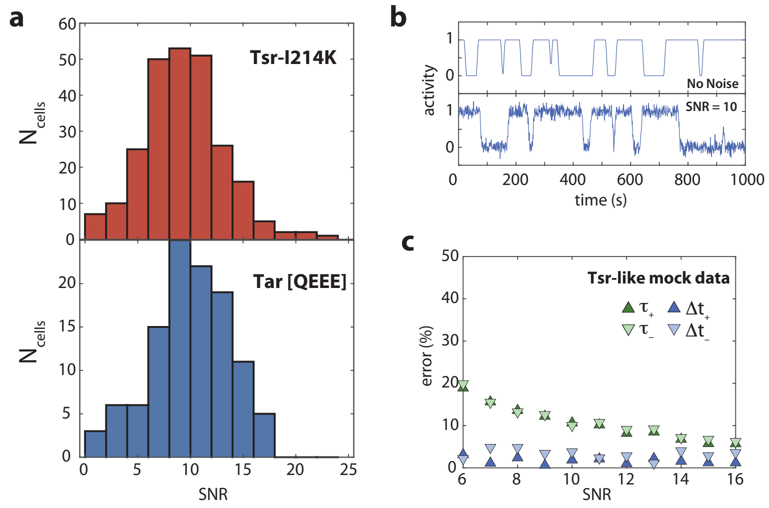


Figure **S4**: **Effect of signal-to-noise ratio on extracting switching parameters.** (**a**) Signal-to-noise ratio (SNR) for two-state switching cells calculated as the maximum FRET response divided by the fluctuations in the FRET signal during attractant response. **(b)** Mock-time series generated by sampling residence times from an exponential distribution and assuming one transition time (top panel) with Gaussian white noise added to simulate experimental noise (bottom panel). **(c)** Relative uncertainties for transition times $\tau$ and residence times $\Delta t$, based on analyzing two state time series with added Gaussian white noise. The simulated time series are based on the average transition and residence times measured for Tsr-I214K.

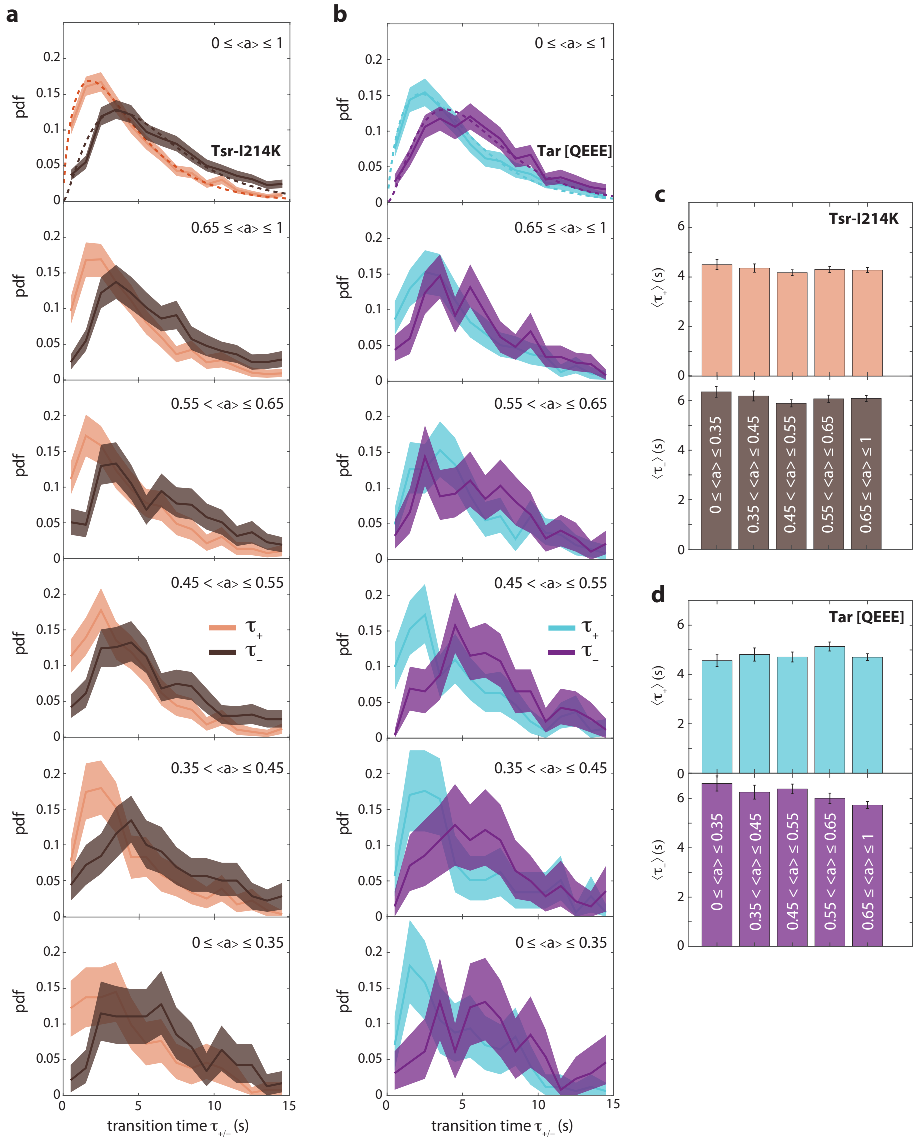


Figure **S5**: **Transition time distributions sorted by cellular activity bias. (a)** Histograms of transition time events across 549 cells expressing Tsr-I214K with each event sorted by the activity bias of the corresponding cell. Shaded areas represent 95% confidence intervals obtained through bootstrap resampling. Dashed lines: gamma distribution fits with the following parameters: upwards transition: $\alpha = 1.72$ and $\beta = 2.5$; downwards transition: $\alpha = 2.45$ and $\beta = 2.48$. **(b)** As in (a), but with 204 cells expressing Tar [QEEE]. Gamma distribution fit parameters: upwards transition: $\alpha = 1.87$ and $\beta = 2.57$; downwards transition: $\alpha = 2.74$ and $\beta = 2.2$. **(c)** Histograms of mean transition time per activity bias for cells expressing Tsr-I214K; (top) mean upwards transition time, (bottom) mean downwards transition time. Error bars represent standard error of the mean. Upwards and downwards transition times are independent of the activity bias. **(d)** As in (c), but for cells expressing Tar [QEEE].

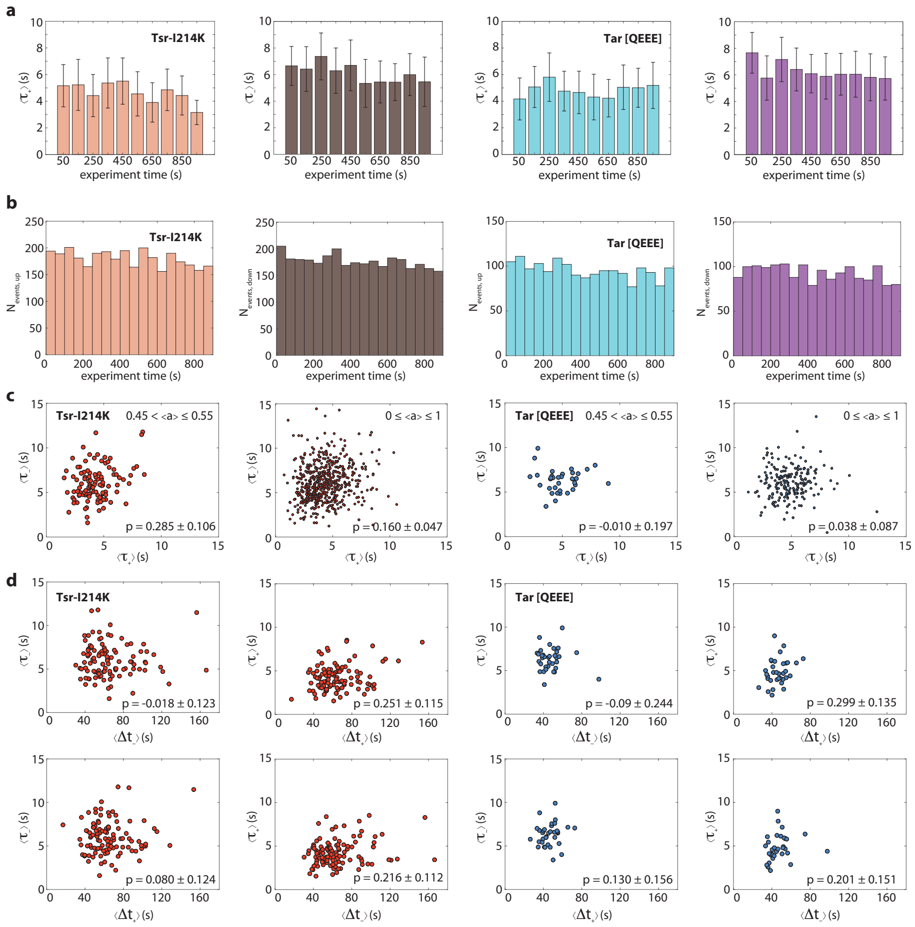


Figure **S6**: **Distributions of incidence times of switching events and correlations between transition and residence times.** **(a)** Distribution of mean upwards (orange) and downwards (brown) transition times with respect to the incidence time of each transition for cells expressing Tsr-I214K; turquoise and purple: same as previous, but for cells expressing Tar [QEEE]. Error bars represent standard deviation. Mean transition times are constant throughout the duration of each experiment. **(b)** Number of upwards (orange) and downwards (brown) events with respect to the incidence time of each transition for cells expressing Tsr-I214K; turquoise and purple: same as previous, but for cells expressing Tar [QEEE]. The frequency of transitions is constant throughout the duration of each experiment. **(c)** Correlations between upwards and downwards transition times for cells expressing Tsr-I214K (red points) and cells expressing Tar [QEEE] (blue points). Error margins in Pearson correlation coefficients are 95% confidence intervals obtained through bootstrap resampling. **(d)** Correlations between transition and and residence times for cells that have an intermediate activity bias and express Tsr-I214K (red points) or Tar [QEEE] (blue points). Error margins in Pearson correlation coefficients are 95% confidence intervals obtained through bootstrap resampling. Only small strength of association is observed between these switching parameters.

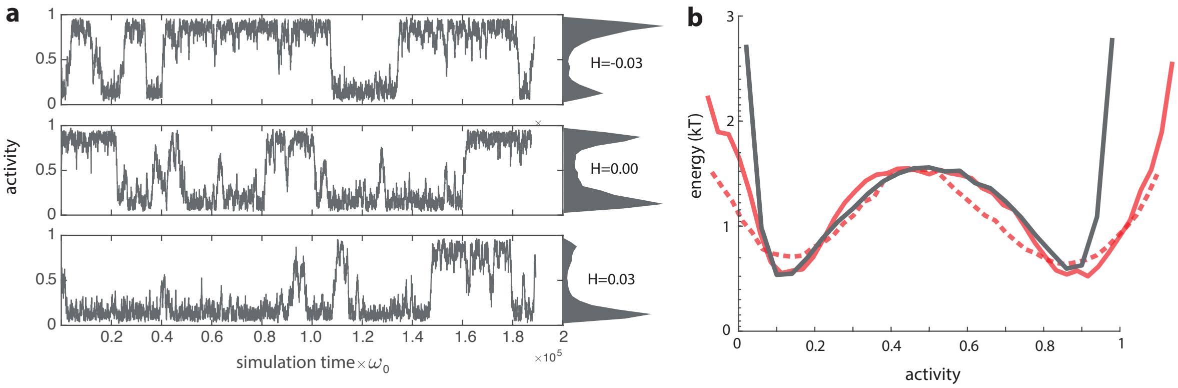


Figure **S7**: **Time series and energy landscape of simulated time series.** **(a)** Example time series and histograms from simulations of the conformational spread model with N=22, coupling energy J=0.475 $k_B T$, and external field as indicated. **(b)** Energy landscape based on 8 simulation runs as shown in panel (a) with H=0, calculated as -log of the histogram (grey). For comparison, the energy landscape based on the experimental FRET time series is superimposed and scaled horizontally and vertically to compare shape, both for the raw time series. The curve from Fig. 2b based on a 7s moving average filter (red, solid) is shown together with the energy landscape based on the raw FRET data (red, dashed).

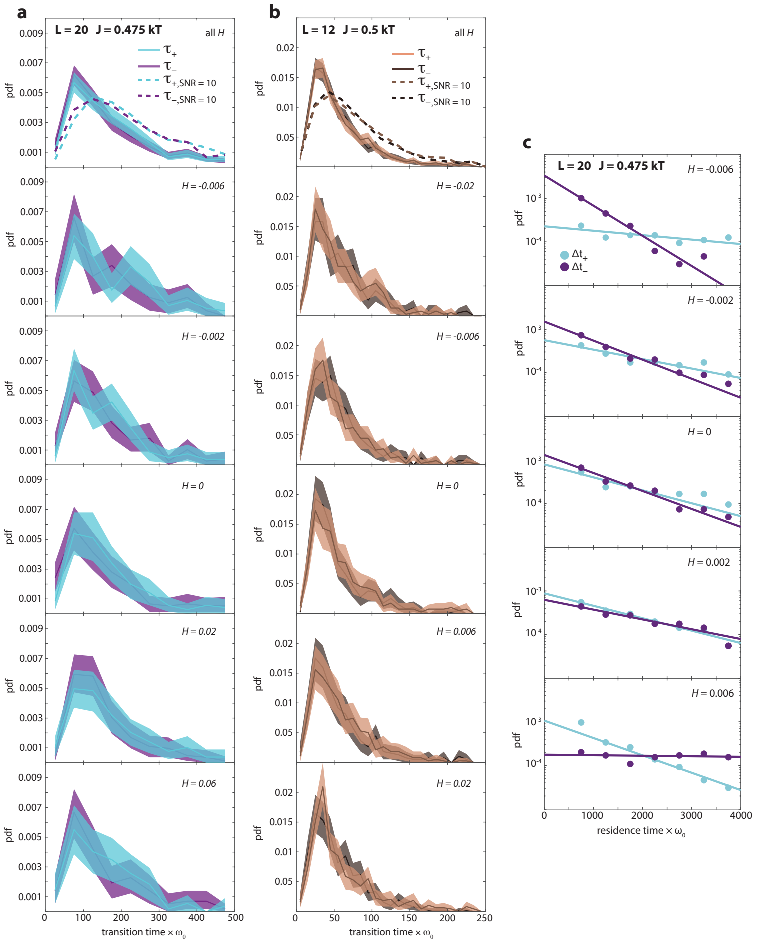


Figure S8: **Transition time distributions obtained from numerical simulations using a conformational spread model.** **(a)** Histograms of transition time events from Ising simulations with a lattice size L of 20 by 20 spins and coupling energy of J = 0.475 $k_BT$. Each event is sorted by the bias $H$ of the external field. Shaded areas represent 95% confidence intervals obtained through bootstrap resampling. $N$ represents number of transition events. **(b)** As in (a), but from Ising simulations with a lattice size L of 12 by 12 spins and coupling energy of J = 0.5 $k_BT$. Similar to the experiments, the average transition times do not depend on the activity bias (here caused by an external field). The Ising lattice sizes and coupling energies were chosen in both cases to recapitulate the ratio of the residence over transition times determined experimentally for Tsr. **(c)** Residence time events from Ising simulations with a lattice size L of 20 by 20 spins and coupling energy of J = 0.475 $k_BT$, together with exponential fits to the data (dashed lines). Each event is sorted by the bias $H$ of the external field.

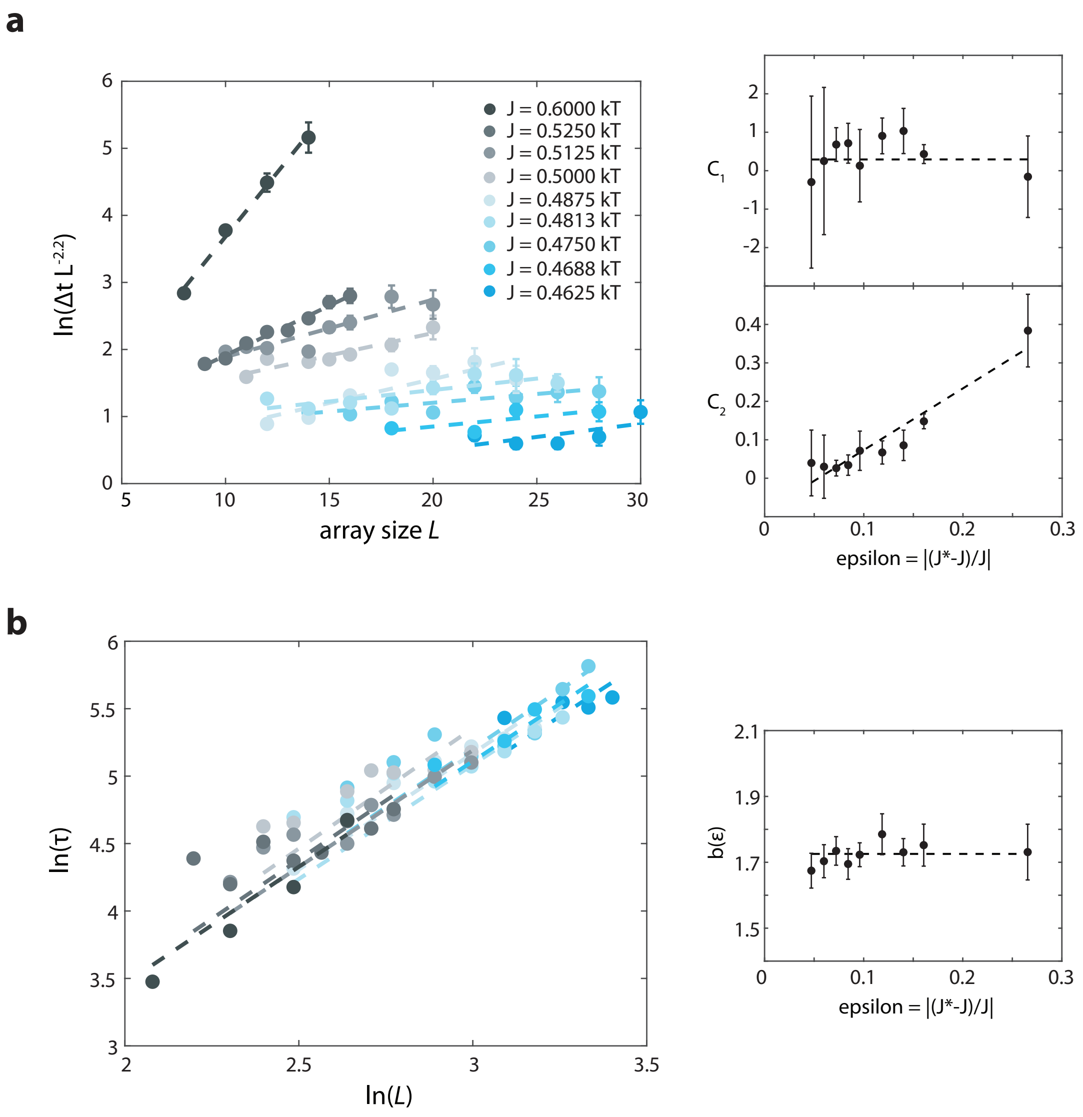


Figure S9: **Mean residence and transition times as a function of lattice size obtained from numerical simulations of a conformational spread model.** **(a)** (Left) mean residence times as a function of array size $L$, scaled according to equation S6. Dashed lines are fits of a function of the form $c_1 + c_2 L$ to the data and colors indicate different coupling energies (the same color scheme is used throughout the figure). Error bars represent standard error of the mean. (Right) scaling of the fitting parameters $c_1$ and $c_2$ as a function of the energy level $\epsilon$. **(b)** (Left) Logarithm of mean transition times as a function of the logarithm of the array size $L$. Dashed lines are fits of the linear form $\ln(\tau) = b(\epsilon)\ln(L)$; (right) Fitted values of $b(\epsilon)$ demonstrated nearly no dependence on $\epsilon$, and hence in our scaling analysis we treated the exponent $b$ as fixed parameter that is independent of $\epsilon$.

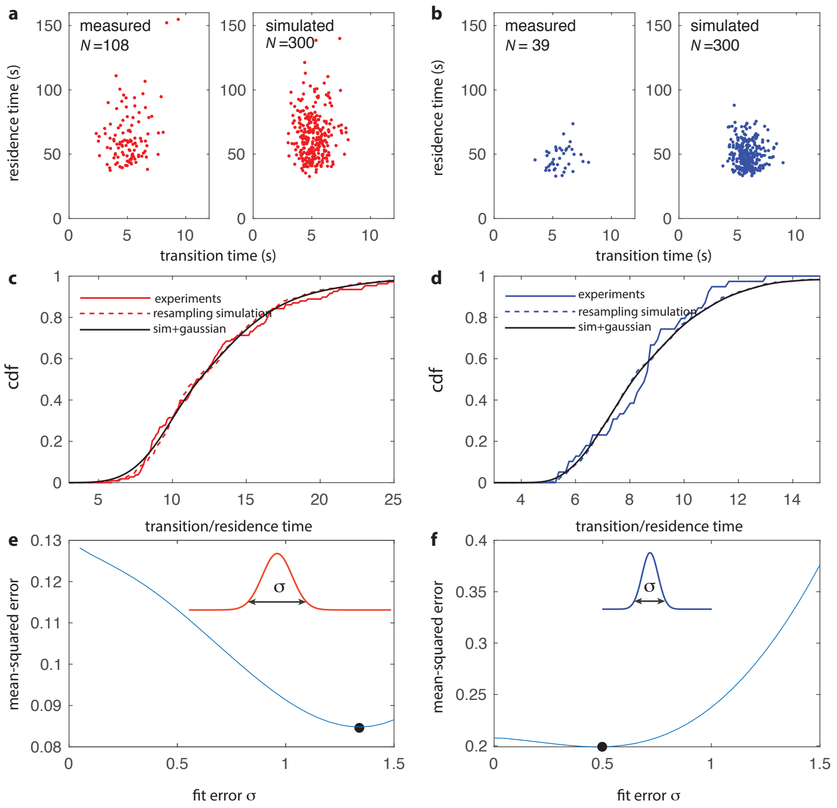


Figure **S10**: **Estimation of the cell-cell variation in transition/residence time ratio based on convolution of true variation with sampling error.** **(a)** Measured transition time vs residence time for (Left) 108 cells expressing Tsr from FRET experiments, with activity $0.45 \leq \langle a \rangle \leq 0.55$, and (Right) for 300 simulated cells. For each simulated cell, residence times and transition times were drawn respectively from an exponential and gamma distribution with parameters extracted from the experiments (Figs. 2 and S5), where the sum of residence times was limited to the experimental duration (1000s). The variation in the simulation represents the sampling error that is observed without variation in parameters between cells. **(b)** As in (a) but for cells expressing Tar. **(c)** The cumulative density function (cdf) for the distribution of transition/residence time ratio per cell in experiments (solid red line) and in the simulated data (dashed line). The simulated cdf (sampling noise) was convolved with a Gaussian pdf (cell-cell variation in ratio) with variance $\sigma^2$, representing true cell-cell variation in timescale ratio, and superimposed on the curve (black line, $\sigma = 1.1$.) **(d)** As in (b) but then for cells expressing Tar ($\sigma = 0.5$). **(e)** Mean squared error (mse) between the convolved cdf (resampling cdf $*$ gaussian pdf) and the experimental cdf, as a function of the standard deviation $\sigma$. The value of $\sigma$ that minimizes the rmse represents the cell-cell variation in timescale ratio that best describes the experimentally observed variation. **(f)** As in (e) but then for Tar.

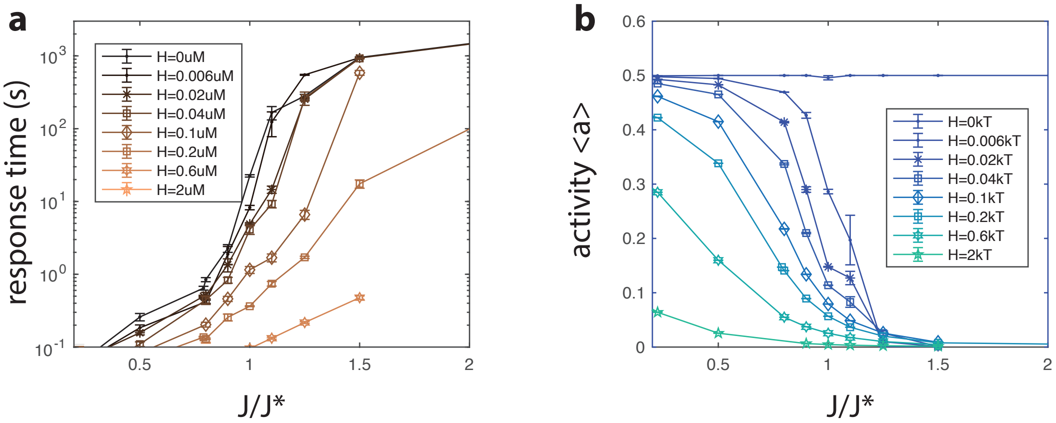


Figure **S11**: **Determining response time and activity level with numerical simulations (a)** The response time as a function of coupling energy $J$ and external field $H$. Response times larger than 100 seconds are lower bound estimates. **(b)** Separate extended time simulations generated dose response curves for different levels of $H$ and $J$. Steady-state activity level $a$ was calculated as the average activity level for times longer than the response time.
